# Cerebellar activity is triggered by reach endpoint during learning of a complex locomotor task

**DOI:** 10.1101/2023.10.10.561690

**Authors:** Andry Andrianarivelo, Heike Stein, Jeremy Gabillet, Clarisse Batifol, Abdelali Jalil, N Alex Cayco Gajic, Michael Graupner

## Abstract

Locomotion in complex environments depends on the precise timing and active control of single paw movements in order to adapt steps to surface structure and coordinate paws. Such motor control crucially depends on the cerebellum, which is thought to support moment-to-moment prediction and correction of paw trajectories. Although cerebellar activity has been linked to limb kinematics on flat surfaces, it remains unknown how cerebellar cortical neurons encode paw movements when gait becomes irregular and the requirements for precise motor control vary dynamically. To address this question, we developed LocoReach: a new task which combines continuous and discrete aspects of motor control by requiring mice to walk on a runged treadmill, where each step involves reaching for the next rung. Over several days of learning, mice became increasingly proficient at LocoReach, so that they made fewer, longer strides with faster swings and fewer missteps. Through real-time optogenetic disruption of cerebellar processing, we shortened the swing of the perturbed paw highlighting the online contribution of lobule simplex to precise limb control the task. We next investigated the role of the cerebellar lobule simplex during LocoReach learning using electrophysiological recordings, with particular focus on molecular layer interneurons (MLIs) that shape the timing and gain of Purkinje cell (PC) output. When analyzing behaviorally-evoked responses in MLIs and PCs, we found sharp changes in activity around paw-specific transitions from swing to stance and vice versa. Cells in lobule simplex showed clear behavioral specificity: most neurons were tuned to swing-stance transitions of the ipsilateral paw, a large proportion encoded transitions of other or even multiple paws. Specifically MLIs exhibited larger amplitude firing-rate changes during longer strides acquired through learning, indicating increased engagement during higher-demand steps. These results show that cerebellar activity is tightly aligned to defined events in the step cycle, providing a mechanism through which cerebellar cortical circuits can contribute to the precise control of paw placement during adaptive locomotion.

## Introduction

Precise control of movements during locomotion is essential for maintaining balance and preventing falls in challenging environments, such as uneven or unstable surfaces or when navigating obstacles. Locomotion in such conditions is inherently discontinuous, with rapidly fluctuating demands on motor precision and timing. The cerebellum plays a key role in meeting these demands by integrating sensorimotor information to predict the consequences of motor commands and adjust movements in real time to keep them accurate and well timed (Morton, 2006). Despite extensive knowledge about cerebellar involvement in rhythmic locomotion on flat surfaces, it is still unclear how cerebellar circuits operate when animals must continuously adapt to unpredictable, irregular biomechanical demands-conditions under which cerebellar predictions and corrections are thought to be most critical.

During skilled movements like reaching to retrieve food pellets, the intermediate cerebellum helps to achieve precision by influencing the kinematics of the reach (Becker and Person, 2019) possibly through prediction of reach kinematics encoded by population activity (Calame et al., 2023). During locomotion, cerebellar activity has been linked to kinematic variables of individual limbs (Becker and Person, 2019; Heiney et al., 2014; Hewitt et al., 2015; Hewitt et al., 2011; Pasalar et al., 2006; Vinueza Veloz et al., 2015), coordination across limbs (Darmohray et al., 2019; Vinueza Veloz et al., 2015), and whole-body coordination (Machado et al., 2015) or state variables (Lanore et al., 2021). For locomotion on non-flat surfaces such as a horizontal ladder, Purkinje cells in the intermediate cerebellum show step-related rhythmic activity (Marple-Horvat and Criado, 1999). Together, these findings suggest that the cerebellum contributes to both discrete (reaching) and continuous (locomotor) aspects of movement, yet they have typically been studied in isolation (except for Marple-Horvat and Criado, 1999). This raises a central question for our study: how does the cerebellar cortical circuit operate when reaching and stepping demands are combined and when movements become irregular and dynamically challenging in terms of paw placement?

Cerebellar motor control relies on signal processing in the cerebellar cortical microcircuit, where sensorimotor information is integrated by Purkinje cells (PCs), which constitute the sole output of the cerebellar cortex. The precision and temporal structure of PC output is tightly regulated by molecular layer interneurons (MLIs), which, together with PCs, receive parallel fiber input and form a powerful feed-forward inhibitory motif (Arlt and Häusser, 2020; Barmack and Yakhnitsa, 2008; Duguid et al., 2012; Eccles et al., 1967; Mittmann et al., 2005). In turn, MLI mediated inhibition controls the timing and tunes the kinematics of motor output (Heiney et al., 2014). Previous *in vivo* work has shown that MLI firing rates increase globally during locomotion compared to rest (Bao et al., 2020; Jelitai et al., 2016; Ozden et al., 2012), but their detailed contributions during the stepping cycle, specific locomotor events and online motor control remain largely unexplored.

PC activity has been extensively examined during locomotion and consistently shown to be modulated by the step cycle (Armstrong and Edgley, 1984; Edgley and Lidierth, 1988; Muzzu et al., 2018; Orlovsky, 1972; Sarnaik and Raman, 2018; Sauerbrei et al., 2015). Notably, these step-cycle studies were conducted exclusively on flat surfaces (except for Marple-Horvat and Criado, 1999), where gait patterns are highly regular and biomechanically stereotyped. In such conditions, cerebellar activity may primarily reflect rhythmic, cyclic components of locomotion. In contrast, the task used here involves an uniformly spaced rung surface that introduces substantial variability in paw placement, and corrective movements. This enriched biomechanical context requires continuous sensorimotor updating, precise prediction of paw trajectories, and rapid adjustments to perturbations – conditions under which cerebellar contributions has been demonstrated (Vinueza Veloz et al., 2015). Such irregular locomotion provides a powerful probe into cerebellar circuit function. Notably, no study to date has examined MLI and PC activity during locomotion in a context that demands continuous prediction and rapid correction, leaving a major gap in our understanding of how feed-forward inhibition shapes PC output during adaptive movement.

To elucidate the role of MLIs and PCs in locomotor control and learning, we investigated their activity during skilled limb movements and locomotor learning in mice using a novel task: LocoReach. This task requires mice to advance across a motorized ladder-like surface, where each step demands precise limb placement and controlled forward extension, analogous to reaching movements. We developed the LocoReach task specifically to introduce structured biomechanical challenges, enabling us to dissect underlying cerebellar computations. Through real-time optogenetic inhibition, we demonstrated the involvement of the cerebellar lobule simplex and specifically MLIs in LocoReach task execution. We next performed single-cell electrophysiological recordings of MLIs and PCs during the LocoReach task. We found that the majority of neurons encode swing and stance phases with precisely timed firing rate changes related to stride events, especially at transitions between swing and stance. We observed that MLI engagement increased linearly with swing length as mice adapted their locomotion. We found that a substantial fraction of MLIs and PCs were selective for transitions of more than one paw, even when using a sparsity-constrained model to control for correlations between different paws. Our results reveal specialized MLI and PC firing patterns during locomotion, with sharp, time-locked modulations around swing-to-stance and stance-to-swing transitions—moments when precise paw placement is required on the runged surface and motor control demands are naturally highest. Our findings shed a light on the contribution of cerebellar activity during locomotor adaptation that could play a central role for sculpting cerebellar output to achieve motor precision and learning.

## Results

### LocoReach task allows for finely resolved paw movement analysis in a complex environment

We developed a motorized treadmill with rungs to study locomotor learning and the control of skilled limb movements. Naïve mice were head-fixed on top of a treadmill with regularly spaced rungs and a constant baseline speed of 7.5 cm/s was imposed during the motorization period. Mice had to rely on whisker-guided walking for LocoReach as the task is implemented in complete darkness (see supplementary video 1). A mirror mounted inside the wheel at 45*^◦^* allowed both side and bottom views to be captured using infrared illumination and a high-speed camera (Fig. 1A). We tracked all four paws (front right - FR, front left - FL, hind right - HR, hind left - HL) in the bottom view with high accuracy (Fig. 1B,C; see Materials and Methods for more details).

**Figure 1:**
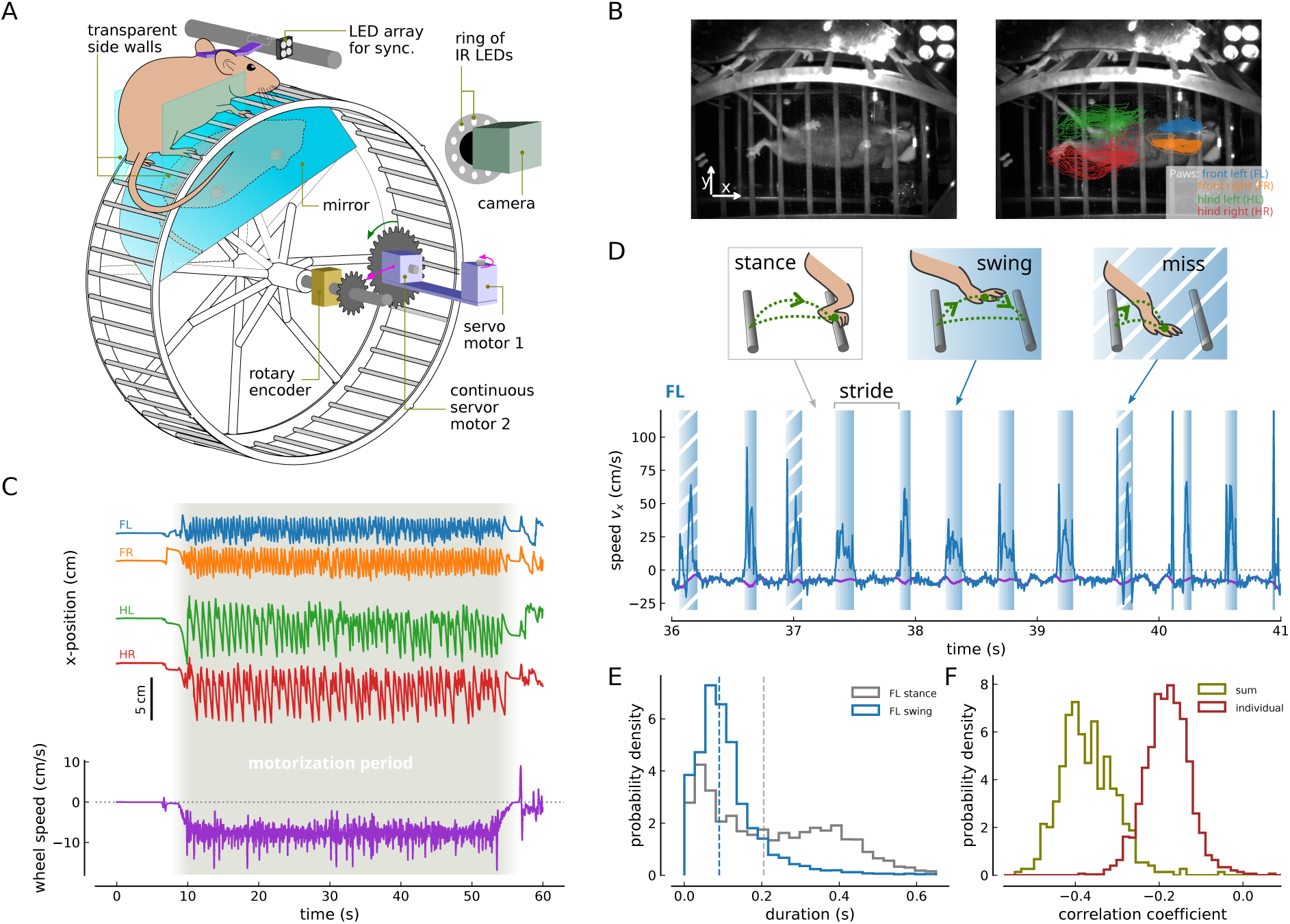
LocoReach is a challenging locomotor task that allows for the analysis of paw specific stepping patterns during locomotion in a complex environment. (**A**) Schematic of the LocoReach setup. Head-fixed mice were placed on top of a runged treadmill with regularly spaced rungs in complete darkness. Mice were illuminated with infrared light. Bottom and side view of the animal were filmed with a high-speed camera (200 fps) through a 45°-inclined mirror under the wheel. During motorization, a servo motor (1) moved in a gear and a continuous servo motor (2) imposed a constant speed. Wheel angluar position was tracked with a rotary encoder. (**B**) Video frames showing a mouse walking on the wheel (left), and with overlayed FL (blue), FR (orange), HL (green) and HR (red) paw trajectories of one recording (right). (**C**) Individual paw x-positions (in the direction of forward locomotion, top) and wheel speed (purple, bottom) vs time during a sample recording. The motorization period (shaded region) started at 7 s with a smooth acceleration, followed by a constant speed period, [10, 52] s, and a smooth deceleration. (**D**) Wheel (purple) and FL paw speed vs time. Swing (changed regions) and stance phases were identified based on the speed difference between the two. Swing and stance phase together constitute the stride of a single paw. Miss steps occurred when the paw did not land directly on a rung and had to correct its position, leading to a multi-phasic paw speed profile (diagonally-striped swings). (**E**) FL swing and stance duration histograms. Dashed lines correspond to median at 0.09 s for swing and 0.205 s for stance. (**F**) Correlation coefficient distributions between wheel and paw speeds. Wheel speed is correlated with the sum of all four relative paw speeds (olive), and the individual relative paw speeds (brown). Distributions in E and F comprise all recordings (*R* = 492) from all mice (*N* = 11).

To classify each paw trajectory time series into swing (paw in the air) and stance (paw resting on rung) phases, we developed a custom routine based on the difference between paw- and wheel speed and taking into account the paw-rung distance (Fig. 1D, see Methods for criteria). Comparable to freely walking mice at slow walking speeds similar to the imposed speed, paws spent larger fractions of the total stride time in the stance compared to swing phase (all mice and FL paw : swing duration [0.055, 0.09, 0.14] s and stance duration [0.075, 0.205, 0.365] s for [25, 50, 75]th percentile) (Fig. 1E, Machado et al., 2015). In contrast to walking on a flat surface however, swing- and stance durations were highly variable due to irregular paw movements on the runged wheel and the paw could sometimes miss the rung at the end of the swing phase, both effects owing to the difficulty of the task because of the challenging surface. Paw position corrections in particular lead to short stances (*<* 0.1 s) and swings (*<* 0.05 s) (Fig. 1E). Miss steps were characterized by multiphasic paw speed profiles (swings with diagonal stripes in Fig. 1D; Fig. 8A) from the slow-down at the expected paw landing time and the acceleration of the subsequent corrective movement. The miss steps give us a means to quantify paw reaching precision during learning. The imposed wheel speed was not rigid but was modulated by the movement and the force exerted by the animal (Fig. 1C). In particular, swings of all four paws are accompanied by a backward acceleration (in negative x-direction) of the wheel. During forward paw advancement, the animal applies force in the same direction as the wheel’s rotation. This force opposes the animal’s own propulsive force, causing the wheel to accelerate in the backward direction. Because the forward paw direction is defined as the positive x-axis in our coordinate system, this backward acceleration appears as a negative deflection in the wheel-speed trace (Fig. 1C,D,F). Consequently, the wheel speed shows a stronger negative correlation with the summed speeds of all four paws than with any individual paw (Fig. 1F). Variations in the imposed wheel speed therefore reflect the animal’s overall movement state: forward movements of all four paws collectively reduce the wheel speed.

LocoReach was inspired by a task for which the cerebellar involvement has been demonstrated (Vinueza Veloz et al., 2015), while additionally providing access to individual paw trajectories allowing for highly refined behavioral analysis of locomotor dynamics and precision. The head-fixed positioning of mice furthermore opened the possibility for neural recordings during task execution (see below). Together, LocoReach promises novel insights into the intricate interplay between locomotor behavior in complex environments and brain activity.

### Mice refine their step kinematics and paw coordination over learning

To assess locomotor learning on the LocoReach apparatus, naïve, adult mice underwent single daily sessions of multiple recording trials for 8 to 10 days. Each recording trial lasted 1 minute and was followed by a pause of at least 1 minute before the next trial. By imposing a constant walking speed over multiple trials and sessions, we aimed to quantify changes in locomotor parameters and precision for otherwise constant task conditions. Mice were habituated to the setup and the head-fixed position without exposure to the rungs prior to LocoReach learning. Mice could walk freely and self-paced outside the motorization period.

First, we asked how single paw behavior changed over the course of the learning protocol and found that mice showed significant changes in multiple walking parameters across sessions and trials. Specifically, the stride number (number of swings) across trials and across sessions decreased significantly (Fig.2A), which occurred in parallel with an increase in the swing length across trials and sessions (Fig.2B). Resolving swing length for individual paws showed that execution of longer strides occurs for all paws across trials and sessions (Fig. 2C). Fewer and longer strides allowed mice to cover the same distance later during trials as well as sessions.

**Figure 2:**
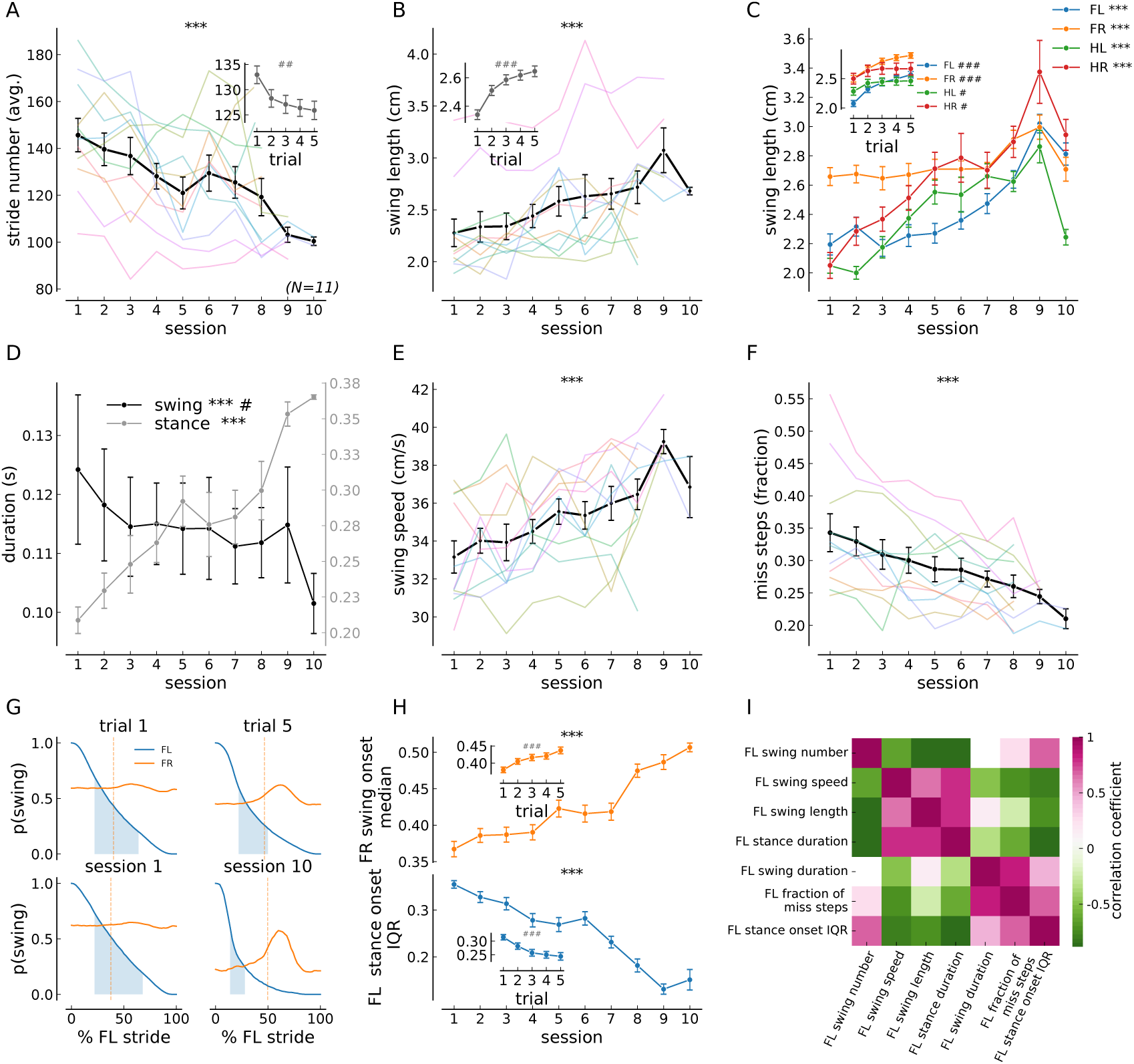
Behavioral learning across trials and sessions. Means of single–paw behavioral measures are shown across sessions; when the trial effect is significant, inset plots show averages across trials (averaged across sessions). Session data are averages over trials within each session for each animal; trial data are averages over all sessions for each animal. Abbreviations: FL, front left; FR, front right; HL, hind left; HR, hind right. (**A**) Stride number (all paws; session effect: *C* = −3.831, *p <* 0.0001; trial effect: *C* = −1.011, *p* = 0.002). (**B**) Swing length (all paws; session effect: *C* = 0.069, *p <* 0.0001; trial effect: *C* = 0.054, *p <* 0.0001). (**C**) Swing length for individual paws (FL: session effect *C* = 0.061, *p <* 0.0001; trial effect *C* = 0.097, *p <* 0.0001; FR: session effect *C* = 0.0240, *p <* 0.0001; trial effect *C* = 0.066, *p <* 0.0001; HL: session effect *C* = 0.088, *p <* 0.0001; trial effect *C* = 0.020, *p* = 0.042; HR: session effect *C* = 0.104, *p <* 0.0001; trial effect *C* = 0.032, *p* = 0.012). (**D**) Swing and stance duration (all paws; swing—session effect: *C* = −0.001, *p <* 0.0001; trial effect: *C* = 0.001, *p* = 0.015; stance—session effect: *C* = 0.013, *p <* 0.0001; trial effect: *C* = 0.001, *p* = 0.100). (**E**) Swing speed (all paws; session effect: *C* = 0.550, *p <* 0.0001; trial effect: *C* = 0.047, *p* = 0.421). (**F**) Fraction of miss steps (all paws; session effect: *C* = −0.012, *p <* 0.0001; trial effect: *C* = −0.001, *p* = 0.328). (**G**) Average Hildebrand plots aligned to FL swing onset, showing the probability of swing for FL (blue) and FR (orange) during trial 1 (upper left), trial 5 (upper right), session 1 (lower left), and the last training session (lower right). The blue shaded area indicates the FL stance-onset interquartile range (IQR), and the vertical orange line marks the median FR swing-onset time. (**H**) Median FR swing-onset time relative to FL swing onset (top; session effect: *C* = 0.013, *p <* 0.0001; trial effect: *C* = 0.012, *p <* 0.0001) and FL stance-onset IQR relative to FL swing onset (bottom; session effect: *C* = −0.021, *p <* 0.0001; trial effect: *C* = −0.014, *p <* 0.0001). Averages across animals are shown over sessions and over trials. (**I**) Pearson correlation matrix between behavioral parameters for the FL paw (example). Colors denote correlation coefficients with *p <* 0.05 (*N* = 11). In (A)–(F) and (H), data are mean ± SEM; black lines show group averages and pale colored lines show individual animals. Statistical significance was assessed with a linear mixed model; regression coefficients (*C*) and *p*-values are reported. *Symbols:* \**p <* 0.05, \*\**p <* 0.01, \*\*\**p <* 0.0001 for session effects; #*p <* 0.05, ##*p <* 0.01, ###*p <* 0.0001 for trial effects.

The stride frequency reduction across sessions was achieved mostly by increasing the duration of the stance phase (Fig. 2D), consistent with what has been reported for walking on flat surfaces for mice and flies (Gonçalves et al., 2022). Swing duration decreased slightly across sessions (Fig. 2D) while swing speed increased to execute the longer strides (Fig. 2E). Also the precision of the strides increased across sessions as quantified by the decrease in the fraction of miss steps (Fig. 2F). Together, we see fewer and longer strides, longer stride durations, and less miss steps demonstrating that head-fixed mice walking on the treadmill with rungs exhibit the same improvements in locomotor performance as freely walking mice on the Erasmus Ladder (Vinueza Veloz et al., 2015).

We next asked whether regularity of single paw movements and inter-paw coordination changed during the protocol by computing the relative timing of front paw swing phases. Using the FL paw as an exemplary reference, two changes were apparent across trials and across sessions: (i) the swing duration of the FL paw became less variable, and (ii) the swing phase of the front right paw started more consistently at 50 % of the front right stride cycle later in training, as illustrated in the Hildebrand plots in Fig. 2G. The regularity of the FL swing phase was quantified by the interquartile range of the stance onset time which decreases over sessions (Fig. 2G,H). The entrainment of the FR paw with the FL can be observed in the median swing-onset time which indicates that FR and FL develop a 50 % phase offset over learning as evidence of more coordinated movements (Fig. 2H). The same improvements in inter-paw coordination and increase in regularity of individual paw cycles can be observed when using the FR paw as reference (Fig. A.1).

In summary, the improvements in motor performance, precision, regularity and coordination indicate that mice were adapting to the wheel speed and rung spacing and became more precise in their movements. This gradual progression reflects substantial initial unfamiliarity and a relatively slow learning rate across days—features that motivate our use of the term ‘complex environment’ or ‘complex task’. The changes in the walking parameters are highly interdependent, *e.g.*, swing speed is positively correlated with stride length and stance duration, while it is negatively correlated with stride number and fraction of miss steps (Fig. 2I; A.1C). The correlations between front-paw kinematic variables underscore that locomotion is a coordinated process in which the movements of individual paws are tightly interdependent, together reflecting improvements in overall gait.

### Real-time optogenetic interference impacts early locomotor performance

Several studies have implicated the intermediate cerebellum in locomotor adaptation (Darmohray et al., 2019) and skilled reaching (Becker and Person, 2019). However, direct perturbations of Purkinje cell activity produce pronounced motor deficits and gross alterations of gait (Darmohray et al., 2019; Vinueza Veloz et al., 2015), which can mask more subtle, timing-dependent circuit computations. To probe cerebellar contributions at finer temporal resolution, we instead targeted MLIs using selective and temporally restricted optogenetic inhibition. Because MLIs shape the timing and structure of Purkinje cell inhibition without directly silencing cerebellar output, this approach allows disruption of intracortical processing while largely preserving overall firing levels. We therefore applied real-time, phase-locked optogenetic inhibition of MLIs in the simplex during the swing phase to causally test their contribution to precise limb control during LocoReach.

Because swing onset and termination are critical for precise paw placement, we inhibited MLI activity selectively during the swing phase of the front left (FL) paw (Fig. 3A). For this purpose, we expressed the blue-light–sensitive inhibitory opsin GtACR2 (Forli et al., 2018; Govorunova et al., 2015) in cerebellar MLIs with high specificity and efficacy (expression in MLIs: 97.47 ± 1.25 % and PCs: 2.53 ± 1.25 % [mean ± SEM]; Fig. 3B; Fig. A.2). Activation of GtACR2 produces rapid and reliable inhibition of MLI activity, with kinetics well matched to the timescale of stepping behavior (Fig. 3C). Using the speed of the FL paw, which was evaluated in real time, blue light was delivered specifically during the FL swing phase with short temporal delay and good selectivity (Fig. 3A,D,E). Opsin-expressing animals exhibited significantly shorter FL paw swing lengths during the first three sessions compared to the control group (exposed to the same manipulations but expressing tdTomato, Fig. 3F), while the swing lengths of the other four paws remained unchanged (Fig. 3F). No differences emerged in the fraction of miss-steps between opsin- and control mice (Fig. 3G). Notably, the FR paw stance onset time IQR was increased in perturbed animals, again lasting for the initial 3 sessions (Fig. 3H). This result points to a disrupted coordination between FL and FR paw.

**Figure 3:**
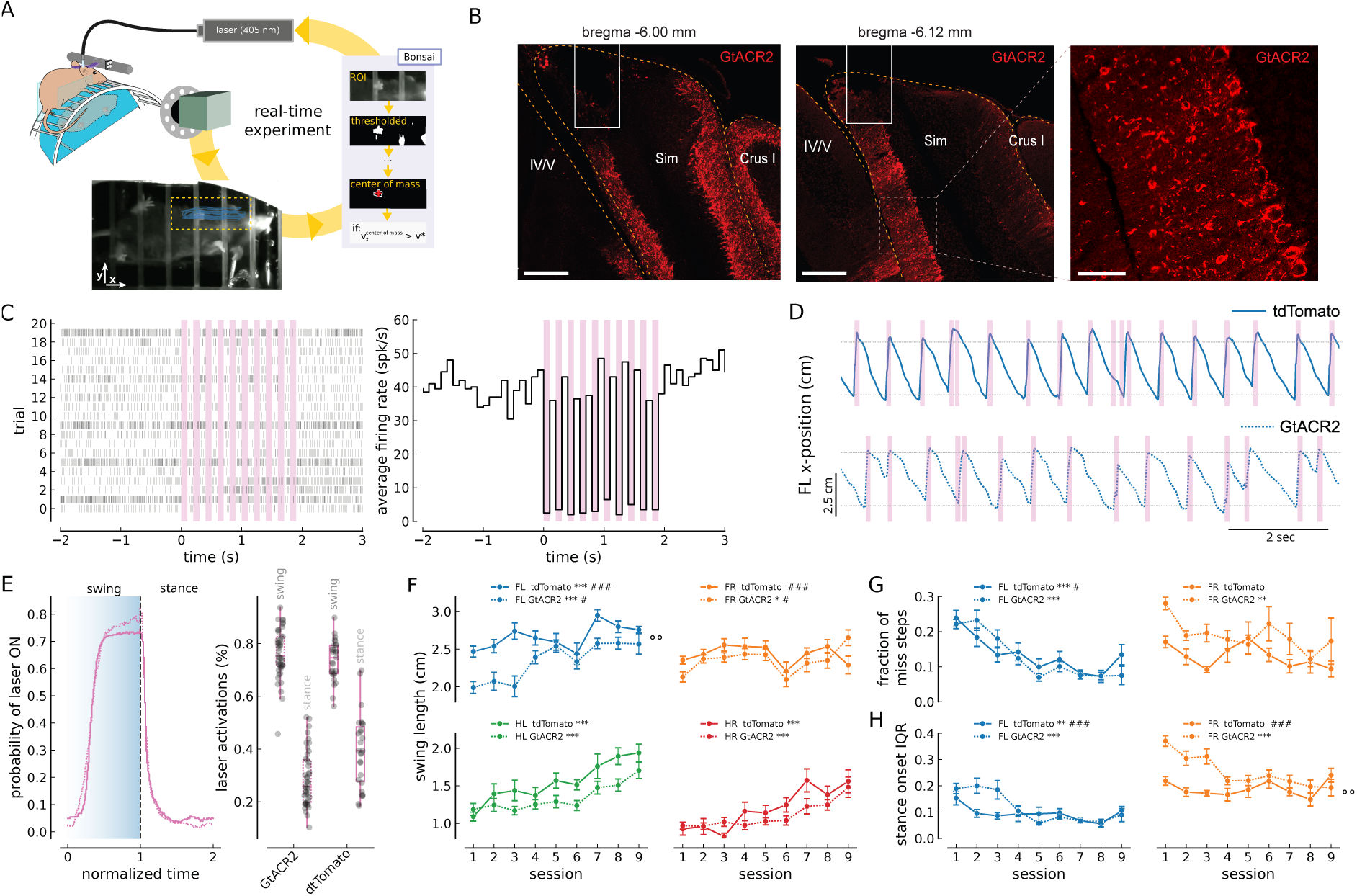
Real-time inactivation of MLIs in lobulus simplex affects swing length and paw coordination. (**A**) Schematic of the real-time inactivation experiment. The video stream was processed online in Bonsai (Lopes et al., 2015) to extract the front-left (FL) paw center-of-mass position and speed. When paw speed exceeded a threshold tuned to detect swing onset, the laser was triggered. (**B**) Photomicrographs showing GtACR2–FusionRed expression in coronal cerebellar sections (scale bar: 200 *µ*m) from two mice (left, middle) and high-magnification image from the left panel Lobulus Simplex (right, scale bar: 50 *µ*m). White squares outline optic-fiber implants. Lobulus simplex (Sim), lobules IV/V of the vermis, and Crus I are indicated. (**C**) Spiking activity from cell-attached recording of an MLI expressing GtACR2–FusionRed during optical stimulation. Laser activation starts at *t* = 0 s, lasts for 100 ms and is repeated 10 times at a frequency of 5 Hz (pink bars). Left panels shows individual trials from 20 repetitions of the optical stimulation. Right panel shows the PSTH of the example neuron. (**D**) FL paw *x*-position during a sample period of a single trial for a tdTomato animal (top, solid line) and a GtACR2-injected animal (bottom, dashed line). Swing-start–triggered blue-laser activations (80 ms, 30 mW) are shown as pink bars. Horizontal gray bars delimit a 3 cm range in both conditions for comparison. (**E**) Quantification of laser activation across the stride cycle. Left: probability that the laser is ON over a full cycle, where x-ranges [0, 1] is normalized swing and [1, 2] is normalized stance. Curves are session averages for the same animals shown in (D) (tdTomato, solid; GtACR2, dashed). Right: probability that laser activation *starts* during swing or stance across all animals and sessions, reported as [25th, 50th, 75th] percentiles—GtACR2: swing [0.714, 0.753, 0.836], stance [0.197, 0.258, 0.359]; tdTomato: swing [0.686, 0.746, 0.793], stance [0.277, 0.393, 0.484]. (**F**) Paw-resolved swing length for tdTomato (solid, *N* = 3) and GtACR2 (dashed, *N* = 5) animals: FL (blue; tdTomato, session effect *C* = 0.042, *p <* 0.0001; trial effect *C* = 0.101, *p <* 0.0001; GtACR2, session effect *C* = 0.082, *Z* = 6.546, *p <* 0.0001; trial effect *C* = 0.062, *p* = 0.012; treatment effect *C* = 0.434, *p* = 0.009), FR (orange; tdTomato, session effect *C* = −0.003, *p* = 0.752; trial effect *C* = 0.081, *p <* 0.0001; GtACR2, session effect *C* = 0.026, *p* = 0.018; trial effect *C* = 0.049, *p* = 0.022; treatment effect *C* = 0.137, *p* = 0.412), HL (green; tdTomato, session effect *C* = 0.091, *p <* 0.0001; trial effect *C* = 0.005, *p* = 0.759; GtACR2, session effect *C* = 0.044, *p <* 0.0001; trial effect *C* = −0.009, *p* = 0.435; treatment effect *C* = −0.053, *Z* = −0.260, *p* = 0.839), HR (red; tdTomato, session effect *C* = 0.083, *p <* 0.0001; trial effect *C* = −0.018, *p* = 0.321; GtACR2, session effect *C* = 0.047, *p <* 0.0001; trial effect *C* = 0.004, *p* = 0.782; treatment effect *C* = −0.040, *p* = 0.857). (**G**) Fraction of missteps for FL and FR paws: FL (tdTomato, session effect *C* = −0.016, *p <* 0.0001; trial effect *C* = −0.012, *p* = 0.045; GtACR2, session effect *C* = −0.021, *p <* 0.0001; trial effect *C* = −0.001, *p* = 0.810; treatment effect *C* = 0.008, *p* = 0.838), FR (tdTomato, session effect *C* = −0.006, *p* = 0.061; trial effect *C* = −0.011, *p* = 0.088; GtACR2, session effect *C* = −0.012, *p* = 0.004; trial effect *C* = −0.006, *p* = 0.490; treatment effect *C* = −0.074, *p* = 0.230). (**H**) Stance-onset interquartile range (IQR) for FL and FR paws: FL (tdTomato, session effect *C* = −0.007, *p* = 0.004; trial effect *C* = −0.018, *p <* 0.0001; GtACR2, session effect *C* = −0.018, *p <* 0.0001; trial effect *C* = −0.004, *p* = 0.422; treatment effect *C* = −0.043, *Z* = −1.071, *p* = 0.284), FR (tdTomato, session effect *C* = 0.00, *Z* = 0.11, *p* = 0.912; trial effect *C* = −0.024, *p <* 0.0001; GtACR2, session effect *C* = −0.020, *p <* 0.0001; trial effect *C* = −0.008, *p* = 0.166; treatment effect *C* = −0.120, *p* = 0.008). Data are mean ± SEM over nine sessions. Statistical significance was assessed with a linear mixed model; regression coefficients (*C*) and *p*-values are reported for effects across sessions unless otherwise specified. *Symbols:* \**p <* 0.05, \*\**p <* 0.01, \*\*\**p <* 0.0001 (session effects); #*p <* 0.05, ##*p <* 0.01, ###*p <* 0.0001 (trial effects); °*p <* 0.05, °°*p <* 0.01 (treatment effects).

In summary, temporally-specific inhibition of MLIs in lobulus simplex during FL swing phases reduced FL swing length and thereby disrupted front paw coordination. This results demonstrates the implication of molecular layer interneurons in the online control of limb kinematics and coordination during complex locomotion similar to skilled reaches (Becker and Person, 2019; Calame et al., 2023). Interestingly, mice adapted to the predictable optogenetic stimulation after 4 sessions exhibiting no difference to the control group for later sessions (see discussion). Our strategy was designed to specifically interfere with fine temporal processing in the cerebellar cortical microcircuit, revealing that temporally structured inhibition contributes to cerebellum-dependent adaptation.

### Molecular layer interneurons exhibit rich firing rate dynamics during LocoReach task

It has been reported previously that molecular layer interneurons increase their average firing rate during locomotion epochs (Jelitai et al., 2016; Ozden et al., 2012). To study their implication at a step-cycle resolved time-scale during the LocoReach task, we performed loose cell-attached recordings in the left lobulus simplex of the cerebellar cortex providing access to the spike output of single MLIs. These measurements were performed in transgenic animals expressing GCaMP6f in ∼ 50 % of the MLIs and allowing for visually guided electrode approaches in a majority of the recordings (Fig. 4A,B). Some approaches had to be performed blindly due to experimental constraints, which resulted in a reduced number of Purkinje cell recordings.

**Figure 4:**
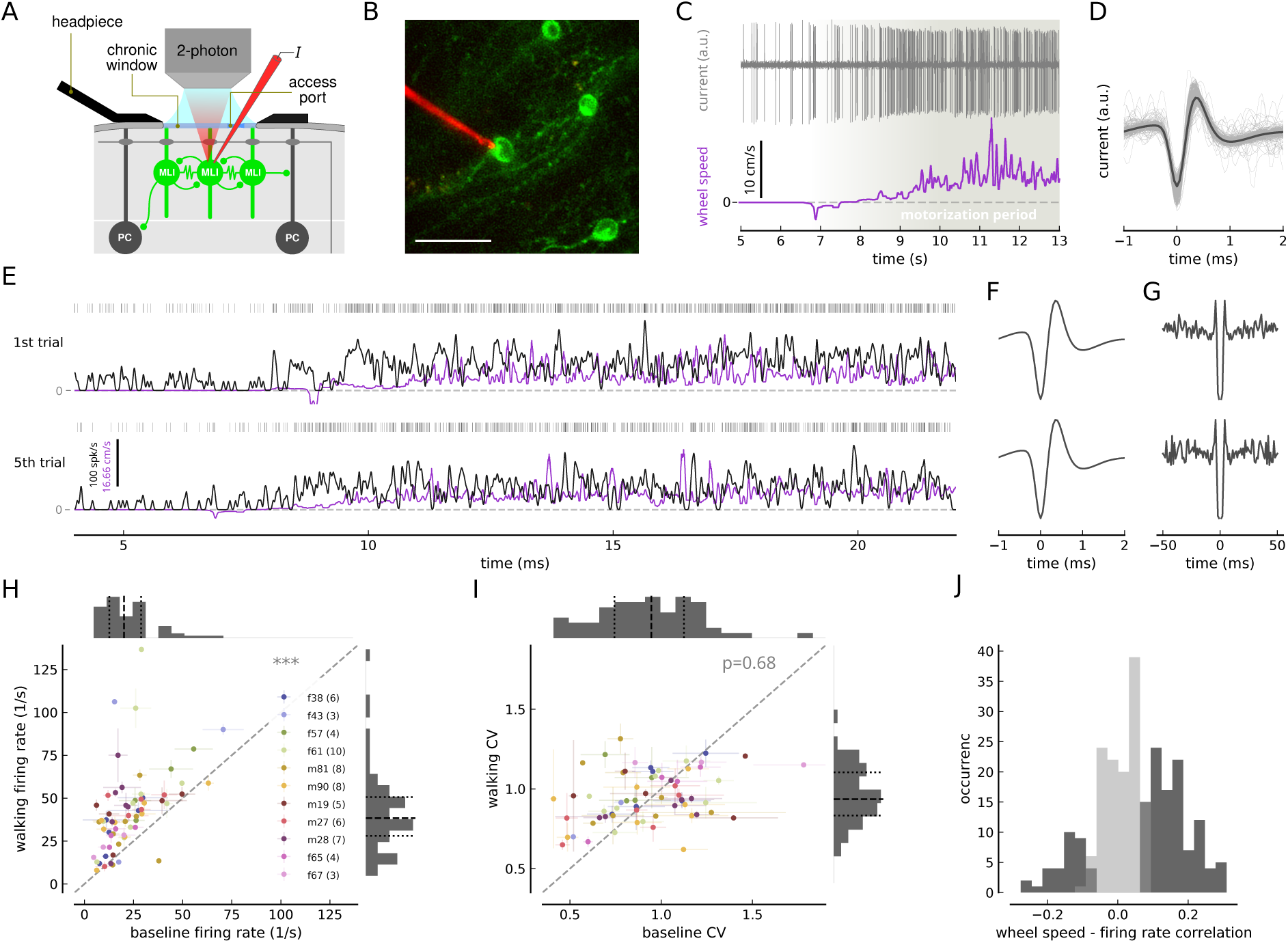
MLIs exhibit firing rate increase and rich dynamics during LocoReach task. (**A**) Schematic of *in vivo* loose cell-attached patch-clamp recordings of MLIs. GCAMP6f-expressing MLIs in transgenic mice were identified using two-photon microscopy through a chronic window and targeted through a silicone access port with a patch pipette loaded with Alexa 594. (**B**) *In vivo* two-photon micrograph of a patch pipette during a recording from a GCaMP6f-expressing MLI (scale bar 25 *µ*m). (**C**) Wheel speed (violet) and example raw, electrophysiological trace (top, grey, high-pass filtered) of a loose-patch recording from an MLI at the initiation of the motorization period (engagement of the gear at 7 s). (**D**) Individual (gray) and average (black) spike waveform of the MLI from (C). (**E**) Wheel speed (violet), instantaneous firing rate (black) and spikes (grey, top) during extracts ([4, 22] s) of consecutive recordings (first and last trial shown) from the same MLI. (**F**) Average waveforms and (**G**) cross-correlograms of an MLI from first (top row) and fifth trial (bottom row, same cell as in (E)). (**H**) Walking period firing rate over baseline period firing rate (paired Student’s t-test: *t* = −7.29, *df* = 63, *p <* 0.0001); and (**I**) walking period CV over baseline period CV (paired Student’s t-test: *t* = −0.42, *df* = 63, *p* = 0.68). Mean and STD are shown across trials for a given cell (panel (H),(I)). Cells recorded from the same animal in same color (see panel (H) where labels correspond to animal ID and total number of recorded cells per animal in parentheses). Top and right histograms show distribution of respective variable together with 25th (dotted), 50th (dashed) and 75th (dotted) percentiles. (**J**) Distribution of walking period correlation between firing rate and wheel speed (*N* = 267 for all recorded trials). Significant correlations (*p <* 0.05) shown in dark gray, and non-significant correlations in light gray

Despite the fact that animals were awake and actively behaving, stable recordings at the transition from rest to locomotion (Fig. 4C,D), during the motorization period, and even across multiple trials provided access to the spike output of single MLIs during an entire behavioral session (Fig. 4C-G; total: 64 MLIs in 267 trials from *N* = 11 mice). We recorded from different MLIs across sessions from the same animal resulting in 3 − 10 cells recorded in total per animal during the learning period (Fig. 4H). We further confirmed the MLI identity of the recorded cells and separated putative Purkinje cells through clustering based on the cell’s electrophysiological signature including spike waveform, spike autocorrelogram, existence of complex spikes, firing rate, and spike-count variability (Fig. A.4).

As expected, locomotor activity was accompanied by a significant increase in MLI firing rate (Fig. 4H; baseline FR [12.7, 20.2, 28.8] spk/s, walking FR [28.1, 38.4, 50.6] spk/s for [25, 50, 75]th percentile). The variability of inter-spike intervals was equally high during rest- and locomotor periods (Fig. 4I; baseline CV [0.74, 0.95, 1.12], walking CV [0.83, 0.94, 1.10] for [25, 50, 75]th percentile). Also Purkinje cells moderately increase their simple and complex spike firing rates during the locomotor period (Fig. A.5A,C), and complex spikes occur more irregularly during locomotion (Fig. A.5D). We next asked whether the rich temporal dynamics of the firing rate during the locomotor period was related to measured task variables. As shown above, the wheel speed reflects the overall locomotor state of the animal including contributions from the individual movements of all four paws. Indeed, we found that the firing rate of more than half of the cells is moderately positively or negatively correlated with the wheel speed (Pearson correlation with *p <* 0.05 for 140*/*267 MLI recordings, Fig. 4J; Pearson correlation *p <* 0.05 for 74*/*143 PC recordings, Fig. A.5C for PCs).

### Stride-related firing rate modulations linked to swing- and stance onset

The observation that the wheel speed is correlated with MLI activity of many cells during locomotion (Fig. 4H) raised the question whether the swing-stance cycle of individual paws modulates MLI activity. The cerebellar simplex projects to the anterior interposed nucleus, where cells exhibit reach-endpoint aligned activity (Becker and Person, 2019) and a signature of this could be present upstream in MLIs during the LocoReach task where paw forward movements resemble skilled reaches. To investigate this possibility, we explored peristimulus time histograms aligned to the extracted transition times from stance to swing and vice versa, called swing- and stance onset.

Strikingly, visual inspection of spike times aligned with the paw positions of an example recording revealed a preferred occurrence of spike discharges right before the peak x-position in many instances (example trace in Fig. 5A). This was true in particular for the FL paw and to a lesser extent for the FR paw. The peak x-position of the paws corresponds to the most forward position during the step cycle, that is, the end of the swing phase (steep upward spokes in Fig. 5A) and the beginning of the stance (slower downward strokes in Fig. 5A).

**Figure 5:**
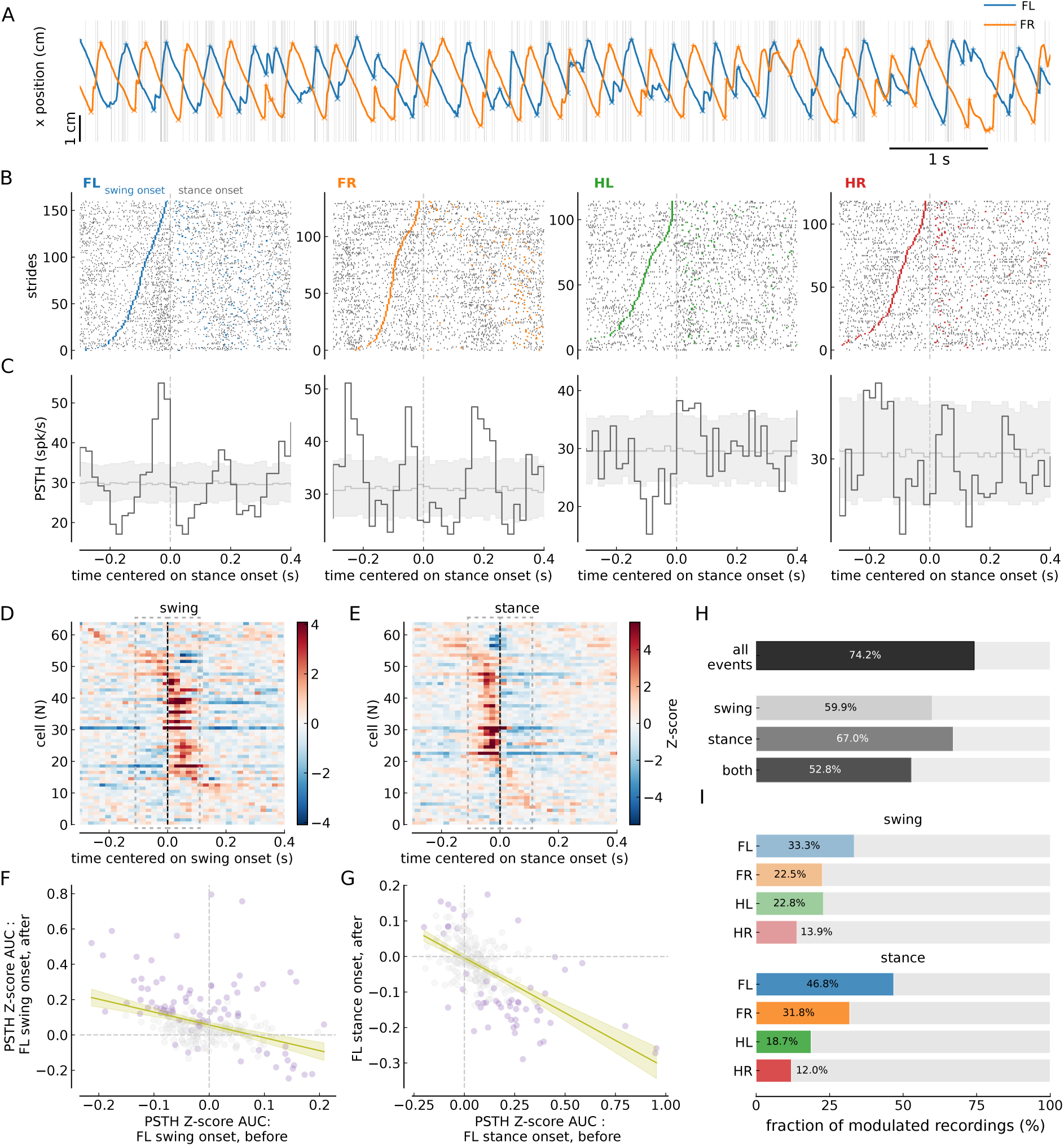
MLI activity modulations at swing- and stance onset. (**A**) Spiking activity of an example MLI (grey) and simultaneously recorded FL (blue) and FR paw (orange) x-positions during a sample period of a single trial. ‘x’ and ‘+’ symbol represent swing onset and and offset, respectively. (**B**) Raster plots showing the occurrence of spikes aligned to stance onset (vertical gray dashed lines) during all strides of each paw (four columns for FL, FR, HL and HR from left), for an example recording of an MLI. Strides are sorted by swing duration. Colored ticks at negative times mark swing onset time for the swing ending at 0 s, and at positive times the swing onset for the subsequent swing. (**C**) Stance-onset aligned PSTHs corresponding to each paw’s raster plot shown in (B). Pale gray lines and shaded area are mean and [5, 95] percentile intervals of 300 stance-onset aligned PSTHs with shuffled spikes times (see Methods for details). (**D**) Swing-onset aligned PSTHs of all MLIs (*N* = 64) for the FL paw. PSTHs are shown as z-scores (see Material and Methods) and ordered according to the occurrence of the positive peak, with early peaks on top and peaks occurring later at the bottom. Each cell’s PSTH is averaged across all recordings of that cell. (**E**) Same as D but for stance-onset aligned PSTHs. (**F**) The area under the z-scored PSTH curve (AUC) before (x-axis) vs after (y-axis) FL swing-onset. Significantly modulated PSTHs before and/or after are shown with purple circles. The criterion for sig. modulation is two consecutive crossings of the [5, 95] percentile interval of shuffled PSTHs during the 100 ms before or after stance onset. The interval of the AUC evaluation (100 ms) in indicated by gray squares (dashed lines) in D and E. A linear regression was performed (Slope = −0.73, Intercept = 0.057, *R*^2^ = 0.139, adjusted *R*^2^ = 0.136, slope p-value *<* 0.001, slope 95% CI: −0.947*…* − 0.509). (**G**) Same as F but for the stance-onset z-scored PSTH curve. Linear regression results : Slope = −0.31, Intercept = −0.004, *R*^2^ = 0.359, adjusted *R*^2^ = 0.357, slope p-value *<* 0.001, slope 95% CI: −0.360*…* − 0.260. (**H**) Bar plots showing the percentage of significantly modulated MLIs for all paws and all events (top), for swing- (2nd row), stance-onset (3 row), or both swing- and stance-onset (bottom row). (**I**) Paw resolved fraction of significantly modulated MLIs for swing- (top) and stance-onset (bottom).

Across stride cycles, FL paw stance-onset alignment revealed a consistent peak activation during the end of the swing phase and a sharp drop in activity at the beginning of the stance phase for this example cell (left panels Fig. 5B,C). FR and HL stance-onset modulations were less pronounced in amplitude and no significant change was observed for the HR paw for the example recording (Fig. 5B,C). For the same recording, spikes were not aligned to stance onset (Fig. A.6A).

We quantified this effect in all MLIs and found that a large majority of cells (74.2 %) show significant firing rate modulations around the step cycle transitions of all four paws, with 59.9 % showing significant changes around swing-onset. 67 % around stance-onset, and 52.8 % with significant changes around both (Fig. 5H). The FL paw stance-onset was the paw specific event which was accompanied most frequently by significant activity changes, *i.e.*, in almost half of the recordings (46.8 %; Fig. 5I) and possibly due to the predominantly ipsilateral control of movement by the cerebellum (Darmohray et al., 2019). Fewer cells exhibited modulations around FR stance-onset (31.8 %), and swing-onset modulations were less common than stance-onset modulations for both paws (FL swing-onset: 33.5 %, FR swing-onset: 22.5 %). Firing rate changes around hind paw transitions were observed but less frequent than with the front paws (Fig. 5I).

Remarkably, firing rate modulations at step cycle events were not temporally symmetric around swing- or stance-onset but instead showed an transition precisely at the event time (Fig. 5D,E). To quantify this asymmetry, we computed the area under the PSTH curve (AUC) in a fixed time window immediately before swing- or stance-onset and plotted it against the AUC in an equally sized window immediately after the transition (Fig. 5F,G). If z-scored firing rate modulations tend to reverse sign at the transition, the resulting cloud of points should align along a line with a negative slope, with the slope direction indicating the predominant change in activity. Consistent with this expectation, linear regression across all MLI recordings revealed a significant negative slope for both swing- and stance-onset transitions(Fig. 5F,G). This indicates that, in most cases, firing rate modulations not only changed significantly at the transition but also flipped sign across the event (Fig. 5F,G; analysis window 100 ms, see Methods for details).

We performed the same swing-cylce-aligned PSTH analysis for the PC population (Fig. A.7). Consistent with previous reports (Armstrong and Edgley, 1984; Edgley and Lidierth, 1988; Muzzu et al., 2018; Orlovsky, 1972; Sarnaik and Raman, 2018; Sauerbrei et al., 2015), PC activity was modulated by the swing–stance cycle in the majority of recorded cells (79 %; Fig. A.7H). Similar to the MLI population, a larger fraction of PCs was modulated around the swing-to-stance transition compared to the stance-to-swing transition (71.3 % vs. 67.1 %; Fig. A.7H), and modulation predominantly involved ipsilateral paw movements (Fig. A.7I). Although we observed a significant sign change in z-scored firing rates around swing–stance transitions (Fig. A.7D-G), the relatively small number of PC recordings limits our ability to distinguish discrete event-related responses from modulation across continuous phases of the stepping cycle.

The anticipatory activity modulation before the end of the stance and swing points to an event-specific role for many MLIs in lobulus simplex. Firing rate modulations occurred more frequently for stance-compared to swing-onset and for the front paws rather than the hind paws (Fig. 5E). Further indication that the observed activity pattern plays a role for the transitions comes from the fact that firing rate modulations before swing- or stance-onset were accompanied by a change in sign right at the beginning of the subsequent phase, *e.g.*, firing rate increases before stance-onset were accompanied by a firing rate decrease precisely timed to stance-onset and vice versa for rate decreases before stance-onset. In summary, we here revealed MLI firing rate modulations linked to events in the step cycle with the highest demands on motor control suggestive of a cerebellar implication for motor output precision and coordination.

### Firing rate modulations are amplified for longer swings

Firing rate modulations linked to specific points in the step cycle could contribute to swing preparation and swing endpoint precision. We showed in Fig. 2 that single paw kinetics and precision as well as inter-paw coordination change over learning. In light of those results, we wondered whether firing rate modulations could be related to variations in swing-specific behavioral variables. We tested this by examining PSTHs across steps and trials for the same MLI, and compared the electrophysiological response with behavior since learning occurred across trials (Fig. 2).

The shape of the PSTH around swing-stance events was remarkably similar across trials for any given cell (Fig. 6A,B for example cells; A.8A; Fig. A.10A,B for example PC). This shows that identical, event-related firing rate modulations are consistently evoked by specific swing-stance transitions. Intriguingly, rate modulations increased significantly when comparing first and last trials of swing- and stance onset modulated cells (FL: Fig. 6C,D; FR: Fig. A.8D,E; but not for PCs Fig. A.10C,D). In other words, if significant firing rate changes existed around swing-stance transitions, their shape was preserved- and the amplitude of these modulations increased across trials. Overall, a majority of MLIs (70.3 %) showed significant rate modulations in at least one of the recorded trials at FL stance onset, and 59.4 % modulations at swing onset.

**Figure 6:**
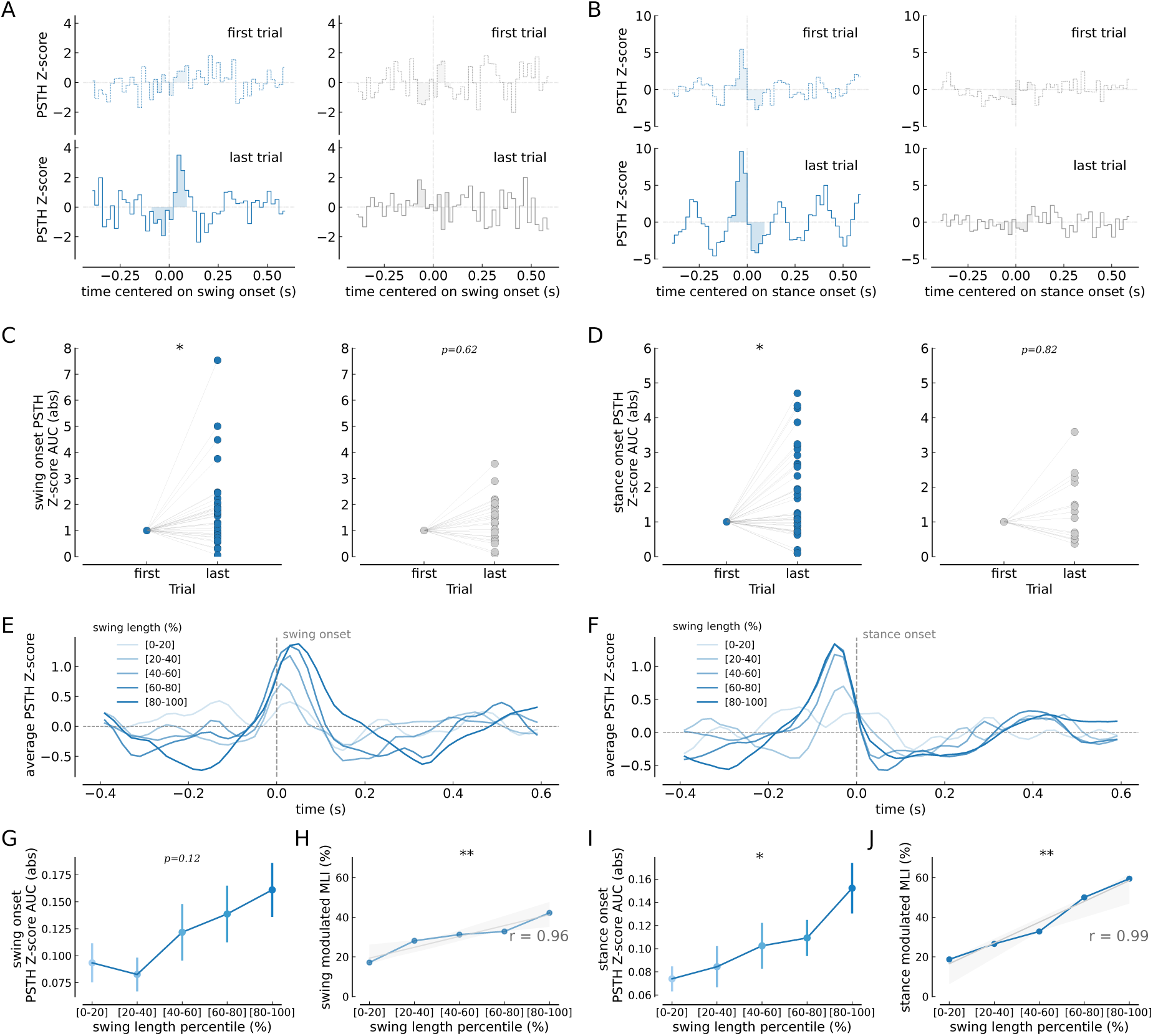
Across-trial FL paw swing- and stance onset-aligned MLI activity correlates with swing length. (**A**) Normalized FL swing onset-aligned PSTHs of a significantly modulated (left) and non-modulated (right) example MLI around swing onset. The z-scored PSTH for first (dashed line, top panel) and last trials (full line, bottom panel) are shown and the area under curve (AUC) is shown as shaded region (analysis window [−100, 100] ms). (**B**) Same as (A) but for a modulated and non-modulated cell around FL stance onset. (**C,D**) The area under the z-scored PSTH curve (AUC) around FL swing onset (C) and FL stance onset (D) is shown for the last trial relative to the first trial of modulated (left panel) and non-modulated (right panel) cells (analysis window [−100, 100] ms; paired Student’s t-test, *p*-values reported in the panels). (**E**) Average FL stance onset-aligned, z-scored PSTH of significantly modulated MLIs for increasing ranges of swing length percentiles. (**F**) Same as (E) but for cells modulated around stance onset (**G,I**) AUC for swing (G, one-way ANOVA, *F* = 1.868, *p* = 0.122) and stance onset (I, one-way ANOVA, *F* = 2.501, *p* = 0.046) of cells modulated for a given range of swing length percentiles. (**H,J**) Fraction of swing onset (H) and stance onset (J) modulated MLIs for ranges of swing length percentiles (Linear regression analysis, correlation coefficient *r*, and the significance of the regression *p* are reported in the panels). Significance is marked according to \**p <* 0.05 and \*\**p <* 0.01.

We next asked whether the event-related firing rate modulations are linked to changes in behavioral parameters. Across trials, mice perform longer steps (Fig. 2B) and firing rate modulations around FL and FR swing- and stance onset increase in magnitude, as shown above (Fig. 6C,D). When combined, we found that activity changes at stance onset are significantly correlated with swing length (Fig. 6I). : the longer the length of the FL swing, the larger the modulation of activity around stance onset (Fig. 6F,I). Furthermore, the longer the FL swing the larger the fraction of cells that are significantly modulated around swing as well as stance onset (for FL: Fig. 6H,J; only for swing onset for FR: Fig. A.8G,I). We note that this difference in modulation occurs for strides within one session depending on their swing length (swing length percentile based analysis of FL: Fig. 6G-J, see methods for details). In contrast, neither the activity modulation magnitude nor the fraction of modulated cells depended on the swing duration across trials (Fig. A.9C-F).

In PCs, the magnitude of activity modulation was not correlated with swing length (Fig. A.10G,I), despite a significant correlation between swing length and the fraction of significantly modulated PCs (Fig. A.10H,J). This dissociation may reflect the greater variability of PC activity profiles around the transition across different swing lengths (Fig. A.10E,F), although we note that the limited sample size may also contribute.

In summary, we found that longer swings imply stronger firing rate modulations for MLIs and a larger fraction of modulated cells overall. On the other hand, swing duration is not correlated with firing rate modulations, consistent with the fact that long lasting steps are not far-reaching steps, *i.e.*, swing length and -duration are not correlated (Fig. 2I). As further reaches imply higher demands on swing endpoint control, a correlation of firing rate and swing length could suggest a role of MLI activity for swing endpoint precision.

Single MLIs get more entrained during learning and encode step cycle events of multiple paws

By analyizing PSTHs of MLIs, we revealed firing rate modulations which occurred at swing-stance (and stance-swing) transitions of specific paws. However, paw movements are periodic and coordinated between different limbs, which could impede a clear attribution of MLI activity modulations to single events. Moreover, as locomotor variables change over the course of a session, it is difficult to discern direct effects of learning on MLI activity from indirect effects, mediated by changes in locomotor parameters (*e.g.* swing length). We therefore devised a multi-step linear regression approach in order to uniquely attribute neural activity changes to single paw events, and to study the dependence of event-related responses on behavioral variables (such as swing length), as well as on learning over trials and across sessions.

We designed a general linear model (GLM) that took into account the discrete transition from stance to swing and vice versa for all four paws to explain single-cell firing rates in MLIs. A sequential L2-L1 regularized regression procedure allowed us to estimate the shape of event kernels, centered around behavioral events, and to add a sparsity bias so that a minimal set of behavioral variables impacts firing rates for each cell (Fig. 7A-C, see Methods for details, Musall et al., 2019). The resulting weighted event kernels captured the characteristic shape of firing rate modulations observed in the PSTH analysis, while controlling for spurious contributions emanating from time-lagged inter-paw correlations (Fig. 7C,D). As before, event kernels point to firing rate increases and decreases before and after swing-stance transitions, often with a sharp transition at the event time (see Fig. 7E for example kernel shapes).

**Figure 7:**
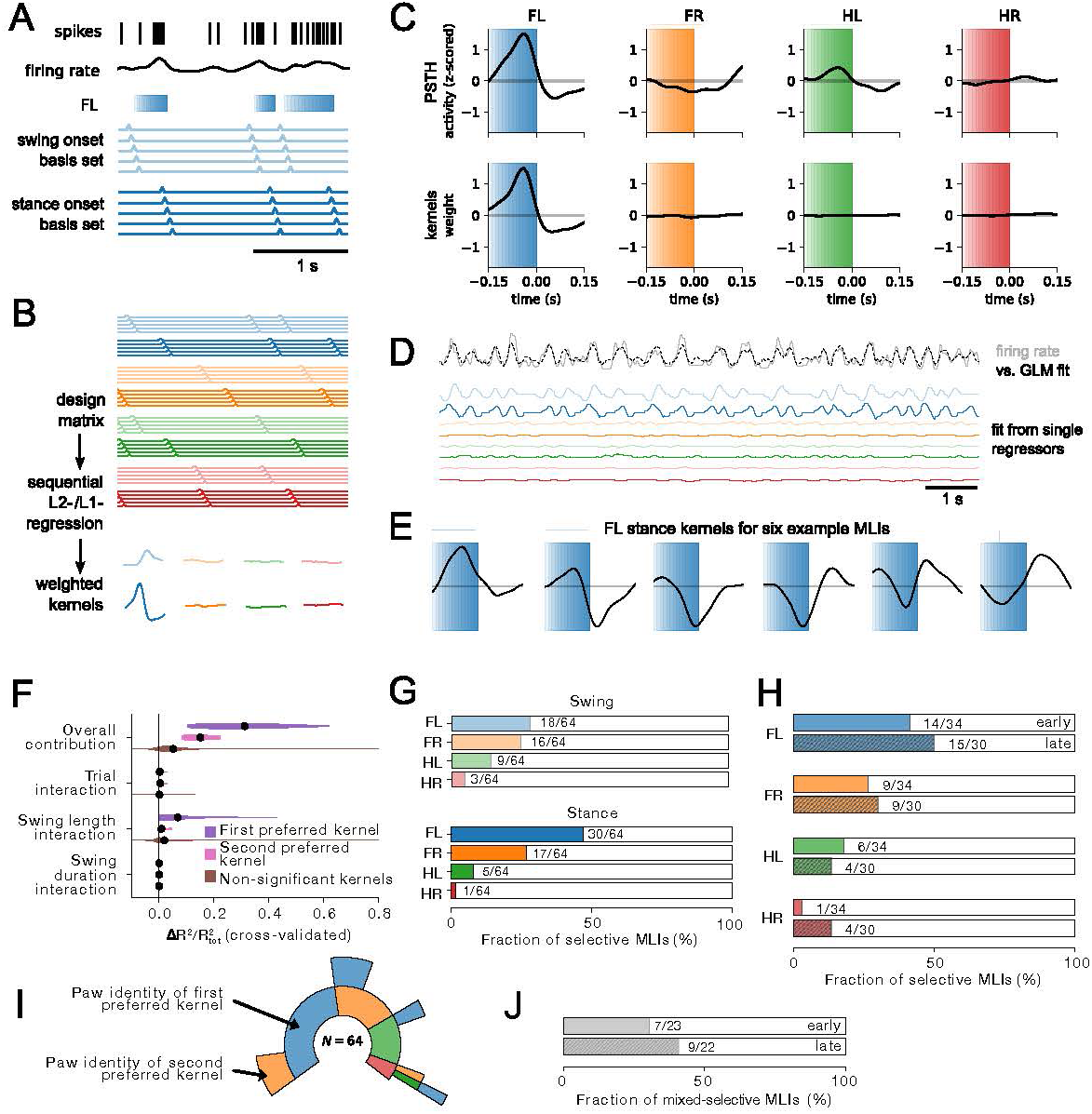
GLM analysis disentangles unique contributions of correlated behavioral events. (**A, B**) Schematic of the two-step GLM analysis of the instantaneous firing rate (black trace). Single-paw swings (graded bars) are encoded as a basis set of time-shifted binary event regressors centered at swing (pale traces) and stance onset (dark traces) (A). Swing- and stance onset basis sets for all four paws form the design matrix for a first, L2-regularized regression yielding event kernels (B) (Musall et al., 2019). Estimated kernels are then convolved with 8 binary event regressors (swing/stance for 4 paws) to form a new design matrix which also includes interaction terms. A second, L1-regularized regression step then provides a sparse representation of the behavioral events impacting firing rate. (**C**) Comparison of the paw specific PSTHs from z-scored firing rates (upper row; see Fig. 5) with the weighted event kernels from the two-step GLM (lower row) for an example cell. (**D**) Snippet from a linear regression example (10 s). Instantaneous firing rate (gray) and fitted firing rate (black) using 8 event regressors and their weighted contributions (colored traces). Paw and event colors as before. (**E**) Example kernels for the FL stance onset event obtained with the two step regression. (**F**) Unique contribution of different regressors (difference in *R*^2^) to overall explained variance 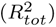. Purple, overall and interaction terms for first preferred event regressor (determined by absolute weight); pink, second preferred regressor; brown, non-significant event regressors. (**G**) Fraction of cells with significant single-variable *R*^2^ for different event types and paws. (**H**) Fraction of cells with significant difference in *R*^2^ (pooling swing and stance regressors) in early (upper, solid bars) and late (lower, striped bars) sessions (see Methods for details). (**I**) Inner circle: 45 out of 64 cells had at least one significant event regressor (swing or stance, difference in *R*^2^), color-coded by paw identity. Second circle, 16 out of 64 cells had a significant second event regressor. (**J**) Fraction of mixed selective cells (relative to cells with at least one significant event regressor), in early (upper) vs. late sessions (lower, striped bar).

We determined the contribution of specific events to single-cell firing rates by measuring the cross-validated *R*^2^ of single-variable models (Musall et al., 2019; see Methods). Most cells responded to stance- (30*/*64) and swing (18*/*64) onsets of the FL paw, with fewer responsive to swing/stance events for the FR- or hind paws (Fig. 7G). This finding confirms results from the PSTH analyses, despite the sparsity constraint of the GLM. We next asked whether model performance could be improved by incorporating continuous variables such as paw position and speed. Adding these regressors did not enhance predictive performance: models containing only continuous regressors or only discrete regressors performed comparably to a full model that included all regressors (Fig. A.12B). Importantly, this similarity in performance arises from strong behavioral collinearity, as evidenced by substantial correlations between continuous and discrete regressors (Fig. A.12A). Because of this collinearity, Δ*R*^2^ comparisons between continuous and discrete regressors provide limited insight into the underlying neural encoding. To more clearly disentangle the role of MLI in encoding locomotion kinematics from event-related encoding of swing and stance onsets, we next focused on missteps, where kinematics and swing/stance events dissociate more clearly (see next section).

We next asked whether learning over sessions was related to stronger encoding of single-paw events. We divided sessions into early (*N* = 34) and late learning sessions (*N* = 30; see Methods for details) for each mouse and compared event encoding across different cells recorded early vs. late in learning. We then asked whether single cells would show increased encoding of transition events. Given that a more coordinated gait late in learning (Fig. 2G,H) might spuriously lead to an increased number of significant events per cell, we measured the unique contribution of an event per session as the difference in *R*^2^ between the full model and a model without the respective event regressor (Fig. A.11C). Even after this control, we found a significant increase in swing- and stance-encoding in cerebellar MLIs over sessions, especially for the front paws (Fig. 7H). Interestingly, we also observed that a fraction of cells were uniquely modulated by more than one paw (Fig. 7I). This fraction of mixed selective cells increased over sessions (Fig. 7J).

Finally, we investigated the fractional contribution of each of the discrete behavioral events to the overall explainable cross-validated variability in firing rate. Not surprisingly, removing the preferred event regressor from the linear model most strongly impacted the explained variance (Fig. 7F; 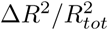 = 0.31 [0.20, 0.31, 0.41], mean, [25, 50 and 75 percentile]). In line with mixed contributions of several paws, removing the second preferred event still reduced explanatory power substantially 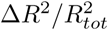 = 0.15 [0.11, 0.14, 0.19]). Using the same approach, we also sought to confirm the differential influence of swing length, swing duration, and trial progression within a session on the strength of event-related responses in MLIs. To this end, we fitted GLMs that contained interaction terms of event regressors with trial numbers, swing length, and swing durations (adding a total of 24 interaction regressors). These extended models accounted for increases or decreases in event-related responses due to learning and behavioral parameters. We found that MLI event-responses were modulated by swing-length (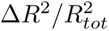 = 0.05 [0.01, 0.030.07]). In contrast, de- or increases in MLI event responses over trials or in relation to swing duration were not significant (Fig. 7F). In other words, progressively longer steps over trials alone explain the increase in modulations in MLI activity (Fig. 7F).

Together, our linear regression analysis demonstrates that most MLIs respond to paw swing- or stance-related events, in accordance with the PSTH analyses. We show that MLIs in left simplex are most responsive to FL stances. Yet, several cells also encode other paw events, and in some cases, cells encode multiple paws at the same time. Over adaptation, we found an increased engagement of MLIs in task execution and an increase in mixed-selective cells. Changes in kernel weights across trials can be explained by taking into account the increase in swing length. Importantly, analyses of the smaller Purkinje cell (PC) dataset revealed qualitatively similar response patterns, with the exception of the correlation with swing length (Fig. A.13). While the limited number of PC recordings precludes strong quantitative conclusions, these observations suggest that key aspects of the event-related structure identified in MLIs may extend to PC. In summary, the newly devised two-step GLM enabled us to capture variations in event-related responses stemming from behavioral factors and learning, singling out swing length as a central variable controlling how strongly swing-stance transition events influence firing rates.

### Miss steps reveal specificity of activity pattern for the swing-stance transition

Both the PSTH and the GLM analyses consistently unveiled activity patterns related to step cycle events. These patterns showed steep changes in activity at the event time, in most cases (Figs. 5D,E and 7E). However, continuous measures describing paw kinematics, such as paw position or speed, also experience abrupt changes at swing-stance transitions. To investigate whether such discontinuities could produce the activity patterns reported here, we conducted a more detailed analysis of the miss steps, which represent 20 to 35 % of all steps (Fig. 2F).

Miss steps are attempted transitions to stance that fail to reach a rung. They are identified by a prolonged reduction in paw speed during the swing phase accompanied by a large distance between the paw and the nearest rung. Miss steps thus provide another reference point in the stepping cycle: the *time of expected stance onset*, when the animal slows its paw and lowers it towards the anticipated rung (Fig. 8A). Failure to contact the rung at this moment forces the animal to execute a corrective movement, advancing or retracting the paw until it successfully contacts a rung.

**Figure 8:**
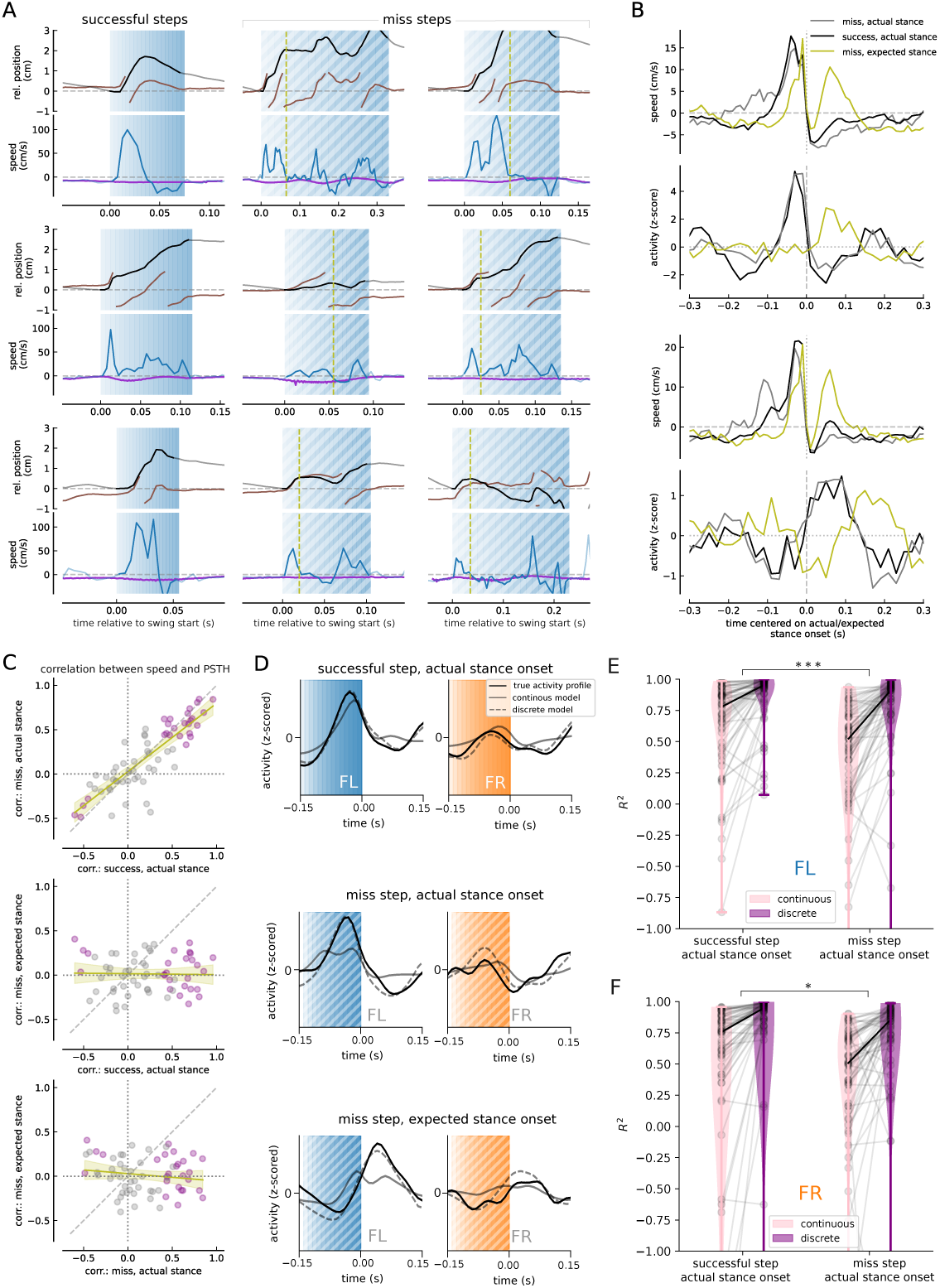
MLI activity of miss steps is modulated around actual and not expected stance onset. (**A**) Focus on kinematics of example swing phases (shaded regions) of the front left paw for successful (left column, blue background) and miss steps (two right columns, blue diagonally-striped background). All curves are aligned to the swing onset at *t* = 0 s. The top row in each panel depicts the x-position of the FL paw (black, normalized to the position at *t* = 0 s) and the signed distance to the closest rung (brown). The lower row depicts the wheel speed (purple) and the speed of the FL paw (blue). The paw position and speed before (*t <* 0 s) and after the swing phase are marked as pale lines. For miss steps (right two columns, diagonally-striped background), the moment of the identified expected stance onset is marked by a vertical line (olive, dashed). All swings are taken from one animal, each row shows steps from a specific session and trial as follows : 1st sess. - 1st trial, 4th sess. - 1st trial, and 8th sess. - 4th trial. (**B**) Comparison of averaged FL paw speed profiles (top row) with the z-scored PSTH for successful and miss steps (bottom row). The two panels (top and bottom) show the averages across the entire recording for two example MLIs. The average FL speed profiles are aligned to the transition from swing to stance for successful steps (black line), miss steps (gray line) and to the time of the expected stance onset for miss steps (olive line). The z-scored PSTHs are separately computed for successful (black) and miss steps and aligned to the same transitions as for the speed profiles. (**C**) Pearson correlation coefficients between paw speed and z-scored PSTH. The correlation coefficients are plotted against each other for different steps and alignments: 1st row shows successful steps vs. miss steps, both aligned to actual stance onset (linear regression results: Slope = −0.77, Intercept = 0.027, *R*^2^ = 0.721, adjusted *R*^2^ = 0.717, slope p-value *<* 0.001, slope 95% CI: 0.65*…* − 0.89), 2nd row for successful steps aligned to stance onset vs. miss steps, algined to expected stance onset (linear regression results: Slope = −0.011, Intercept = 0.016, *R*^2^ = 0, adjusted *R*^2^ = −0.016, slope p-value 0.86, slope 95% CI: −0.14 0.12), 3rd row miss steps aligned to actual stance onset vs. miss steps, aligned to expected stance onset (linear regression results: Slope = −0.084, Intercept = 0.03, *R*^2^ = 0.022, adjusted *R*^2^ = 0.006, slope p-value 0.24, slope 95% CI: −0.23*…*0.06). The correlation coefficient is computed between the corresponding average speed profiles and z-scored PSTH for the interval (−0.4, 0.6) s around the respective alignment point. Purple points indicated that both correlation coefficients (of the two abscissa) are significant (*p <* 0.05). (**D**) Predictions of the GLM model for the swing-stance transition for successful and miss steps. Full black lines: true activity profile; full gray lines: prediction from model with continuous variables as regressors; dashed gray lines: prediction from model with the discrete event kernels as regressors. All curves are averaged over z-scored traces for all respective event types. (**E**) *R*^2^ values in a time window just prior to FL stance onset ([−150, 0] ms) for successful steps (left) and miss steps (right). The *R*^2^ values are compared for predictions between continuous (pink) and discrete regression models (purple). (**F**) Same depiction as in E but for FR paw. For E and F: Permutation test of differences in model performance, true median *R*^2^ vs. median *R*^2^ in 1, 000 permutations; ∗ ∗ ∗*p <* 0.0001 for FL; ∗*p* = 0.04 for FR.

The time of expected stance onset features similar abrupt changes in paw position and speed as the the actual time of stance onset (Fig. 8B) and it is accompanied by the expectation of the animal to enter stance. We therefore asked whether this point in the step cycle evokes similar activity patterns in MLIs as the actual transition to stance. To address this question, we computed PSTH profiles aligned to the expected stance onset for miss steps (Fig. 8B) and compared them to PSTH profiles of successful steps or miss steps aligned to actual stance onset. Remarkably, while average paw speed aligned to expected stance onset shows a bi-phasic profile, MLI activity does not reveal bi-phasic profiles. Rather, the PSTH profile for miss steps appears to be the same as for successful steps, but shifted in time to the actual stance onset time (Fig. 8B). This is in stark contrast to average speed profiles for successful steps, which are mono-phasic, resembling the activity profile exhibited by MLIs with a firing rate increase before stance onset. Consistent with the observation of preserved but shifted MLI activity profiles, we noted that the PSTH of miss steps aligned to actual stance onset resembles the PSTH of successful steps.

We next quantified how closely neural activity and paw speed profiles were linked during successful and miss steps, using alignments to both the actual and the expected stance onset. For each cell, we computed the correlation between its z-scored PSTH and the average paw speed profile. When aligned to the actual stance onset, both successful and miss steps showed consistently strong correlations (Fig. 8C, top row). However, these correlations — whether positive or negative — disappeared when aligning to the expected stance onset (Fig. 8C, middle and bottom rows). This drop reflects the fact that MLI activity no longer tracks paw speed in the period around the expected stance onset.

We further investigated miss steps using the above introduced regression analysis. We observe that around the stance onset of miss steps, discrete models (with event kernel regressors centered on true swing and stance onset) capture the observed MLI activity profile, whereas continuous-valued regression models (with x position and speed for each limb) do not (Fig. 8D). Specifically, in miss steps, MLI activity peaks just before the true stance onset. This pattern is predicted by the stance kernel set. In turn, from x-position and speed regressors, the model predicts a biphasic MLI profile that peaks twice before stance onset (Fig. 8D, second row). We quantified this effect on the group level in an *R*^2^ analysis, where we measured the *R*^2^ only in a time window just prior to stance onset in successful- and miss steps. We expected that the discrete model should outperform the continuous model specifically in miss steps, where the continuous model would falsely predict biphasic increases in MLI activity. Indeed, for front limbs (FL, FR) the discrete model *R*^2^ is comparable to the *R*^2^ of correct steps (Fig. 8E,F, left comparison). In contrast and as expected, the continuous model underperforms specifically for miss step stances (Fig. 8E,F, right comparison).

Together, these analyses demonstrate that MLI firing is not driven by the continuous kinematic changes that accompany step-cycle transitions. Although miss steps contain discontinuities in paw position and speed comparable to those in successful steps, MLIs do not exhibit the biphasic activity that such kinematic profiles would predict. Instead, their responses remain locked to the actual, not the expected, transition to stance. Both PSTH- and regression model-based analyses show that discrete event kernels aligned to true swing–stance boundaries reliably capture MLI activity during miss steps, whereas continuous regressors systematically fail. These findings are consistent with MLIs preferentially encoding discrete step-cycle events over continuous kinematics, supporting the notion that their activity reflects an event-based representation of locomotor state.

## Discussion

To study the role of molecular layer interneurons and Purkinje cells of the cerebellar lobulus simplex in movement precision and locomotor learning, we developed a forced locomotion paradigm of mice walking on a treadmill with rungs allowing to combine detailed analysis of paw movement kinematics with neural activity measurements, called LocoReach. We found that mice are adapting to the wheel speed and the rung distance by improving their locomotor performance over sessions, *e.g.*, they perform longer and more precise steps. We showed for the first time that MLI activity is related to specific events in the gait cycle – transitions from swing to stance and vice versa – most prevalent for the FL paw, located ipsilateral to the recording site, followed by the FR paw and less often observed for the hind paws. Using PSTH and GLM analysis, we demonstrated that activity modulations correlate with swing length. Using real-time optogenetic perturbations during front left swing phases, we support this finding by showing that inhibiting MLIs in lobule simplex output curtails swings in a paw-specific fashion. Later during learning, more cells encode paw events and more cells exhibit mixed-selectivity for more than one paw. Several of our findings reproduce previous results from related tasks. Mice adapt their stepping patterns on runged surfaces by increasing their swing length and reducing the number of miss steps (Fig. 2; Vinueza Veloz et al., 2015). Engagement in locomotion on the runged treadmill elicits a significant increase in MLI activity (Fig. 4H) similar to what has been reported for walking on flat surfaces (Jelitai et al., 2016; Ozden et al., 2012) or walking on a slope (Lyu et al., 2022). In line with the fact that cerebellar control of movement is predominantly ipsilateral (Darmohray et al., 2019; Heffley et al., 2018; Lee et al., 2015), we found that most cells in the left simplex were tuned to ipsilateral front left paw events, also the hind left paw events were more often observed than hind right events (Figs. 5I and 7F,G). Step-related discharges have been reported to be widely distributed in the cerebellar cortex and nuclei during visually-guided ladder walking of cats, including in the intermediate cerebellum, which is the focus of this study, and the interposed nucleus (Armstrong and Marple-Horvat, 1996). Furthermore, the cerebellum influences locomotion largely through projections to midbrain and brainstem premotor nuclei (Liang et al., 2011), several of which preferentially target forelimb motor circuits (Esposito et al., 2014). These anatomical asymmetries likely contribute to the dominance of forepaw tuning relative to hindpaw tuning in our dataset.

Other results are novel and open a new perspective of MLI function in locomotor control. Rather than being linked to continuous kinematic variables of limb movements in PCs (Edgley and Lidierth, 1988; Hewitt et al., 2015; Hewitt et al., 2011; Pasalar et al., 2006) or lick rate in MLIs (Gaffield and Christie, 2017), we showed that MLI activity encodes specific behavioral events suggesting a punctual rather than a continuous task engagement. And MLI engagement occurred at points with heightened demands on motor output control, *i.e.*, the swing start and end points. This aligns with theoretical and empirical evidence that control demands are not uniform across a movement but peak at behaviorally critical phases, such as transitions where spatial and temporal precision are essential (Liu and Todorov, 2007). Beyond ipsilateral movement control, a large number of MLIs responded to contralateral FR and HR events (Figs. 5I and A.8), or showed mixed-selectivity for multiple paws from opposite sides of the body (Fig. 7I). The mixed selectivity for multiple limb movements could allow MLIs to integrate information across cerebellar microzones to coordinate multi-joint dynamics. Modulation of PC activity occurs in relation to the step cycle for walking on flat surfaces (Armstrong and Edgley, 1984; Armstrong et al., 1988; Edgley and Lidierth, 1988; Sauerbrei et al., 2015) or on a ladder (Armstrong and Marple-Horvat, 1996). However, individual cells vary greatly in the phasing of their discharge relative to the step cycle (Armstrong and Marple-Horvat, 1996; Sauerbrei et al., 2015), similarly observed for PCs during skilled reaching (Calame et al., 2023). Our recordings form Purkinje cells support the notion of activity modulations with peaks at continuous phase values (Fig. A.5), but the limited number of cells prevents clear separation of event-related and continuous phase modulation. During walking, population and not single cell activity is at its highest level during the ipsilateral forelimb swing and around the period around footfall (Armstrong and Marple-Horvat, 1996). In contrast, the activity modulation reported here occurs surprisingly consistent across the recorded MLI population linked to the transition events from swing to stance and vice versa, which suggests a more homogeneous MLI population response. This intriguing homogeneity may be attributed to enhanced synchrony within the MLI network, possibly facilitated by gap junctions (Kim et al., 2014; Todd et al., 2025). Consistent with our findings, previous work has shown that MLIs and PCs can display differential firing dynamics. Cerebellar recordings at whisker onset support the notion that that MLIs and PCs can exhibit differential acitivity patterns. While MLIs provide precise, short-latency inhibition for sensory filtering in the input layer, PCs sustain broader, adaptive encoding for motor control (Brown et al., 2024).

Naturalistic motor behaviors are inherently multimodal, requiring the integration of motor commands with multiple streams of sensory information. In the LocoReach task, successful performance depends not only on limb control but also on whisking and somatosensory inputs, which are known to engage cerebellar neurons. Several observations support the relevance of sensory contributions in this task. First, spike rate modulations in the cerebellar simplex have been reported to occur on the time scale of the stepping cycle (Zhai et al., 2024), consistent with the temporal structure of sensory feedback during locomotion. Second, whisking and running are tightly coupled on a step-by-step basis, with whisking rhythms coherent with the stride cycle (Sofroniew et al., 2014). Third, paw kinematics and discrete stepping events account for only a fraction of the variance in MLI activity, suggesting that additional sensorimotor signals contribute to the observed firing patterns. Together, this evidence raises the possibility that MLI activity during LocoReach is partially influenced by whisking-related or other somatosensory signals. However, whisking dynamics are continuous and quasi-periodic, lacking the sharp temporal discontinuities that characterize swing–stance transitions. Even when coupled to locomotion, whisker phase evolves smoothly over time and is therefore unlikely to generate the abrupt, step-locked firing transitions observed in MLIs. Moreover, if whisker–rung contact were partially encoded, one would expect multiphasic activity profiles during longer strides, as multiple rungs are crossed. Such multiphasic patterns are not observed in our data (Fig. 6E,F). Instead, our analysis of MLI activity during miss steps suggests that the activity increase reflects the selection or anticipation of the specific rung targeted for stance, as this transient increase is absent during miss steps prior to the expected stance onset (Fig. 8). Disentangling the respective contributions of limb kinematics and sensory inputs in complex behaviors such as LocoReach will require future experiments. Preliminary data indicating that whisker trimming severely impairs task precision further underscore the importance of somatosensory signals and motivate ongoing work to resolve their specific roles.

Purkinje cells and neurons in the interposed nucleus are modulated during and near the endpoint in a reaching task and this activity is necessary to convey kinematic information as well as to slow the limb near endpoint (Becker and Person, 2019; Calame et al., 2023). The encoding of swing-stance transitions by MLIs in lobulus simplex that we observed could provide a potential source for this endpoint alignment in the cerebellar nuclei. The increased MLI activity precisely timed around stance onset could convey swing endpoint information downstream via graded inhibition of Purkinje cell output. In turn, disinhibition of the cerebellar nuclei near swing termination would then enable encoding of reach endpoint kinematics to adjust endpoint precision. This idea is in line with our finding that longer steps elicit larger activity modulations due to their higher need for kinematic control. Step-length modulated and stance onset locked MLI firing patterns could be an essential part of the cerebellar cortex circuitry that enables encoding of reach-related kinematics in the deep nuclei.

The reduction in ipsilateral swing length induced by real-time optogenetic interference of interneurons provides causal evidence for the involvement of lobule simplex MLIs in controlling paw movements during locomotion. This shortening of the swing phase suggests that perturbing cerebellar cortical activity prematurely truncates forward limb progression. Conceptually, this effect parallels previous findings that limb kinematics scale with activity in the anterior interposed nucleus (IntA), the principal downstream target of lobulus simplex output Becker and Person, 2019. In their study, activating IntA neurons curtailed reaches, whereas in our experiments inhibiting MLIs likely disinhibits Purkinje cells, thereby reducing IntA activity and similarly producing shorter swings. The precise circuit mechanisms explaining why these seemingly opposite IntA activity manipulations yield comparable behavioral outcomes remain to be clarified. In both locomotion and reaching contexts, the cerebellar lobule simplex emerges as a key player in modulating the phase and paw positioning, thereby orchestrating precise and coordinated movements.

Our experiments were designed with unilateral optogenetic interference to achieve a clearer causal link between localized cerebellar activity and ipsilateral forelimb control, while minimizing potential confounds from bilateral suppression. Despite altered locomotor kinetics with optogenetic interference, the fraction of miss-steps remains surprisingly consistent between perturbed and control animals. This observation may unveil underlying compensatory strategies that mitigate the impact of MLI inhibition on task performance, *i.e.*, the animal might recede to smaller steps of reduced demand on position control when perturbed and thereby preserve the fraction of paw end-point misplacements for the FL paw. The trend toward increased missteps in the contralateral paw (Fig. 3H, right panel) and altered HL performance throughout learning (Fig. A.3) further suggest a disruption of interlimb coordination and indicates that compensatory strategies may not fully extend to non-perturbed paws. The disruption in inter-limb coordination might be a direct result of perturbed FL movements, or point to a broader role of the lobule simplex in regulating multi-limb movement patterns. The latter together with our finding of mixed selective cells for multiple paws would provide evidence that the lobule simplex is involved in the online computation and control of limb dynamics critical for coordinating multi-limb movements during locomotion. An important next step will be to decipher cell type specific contributions to the observed locomotor effects.

The predictable effect of the real-time perturbations, always occurring at a fixed delay during the swing phase, got compensated and no difference in locomotor parameters between perturbed and control animals was apparent after a few sessions (Fig. 3D,F). This is reminiscent of the adaptation reported upon repeated mossy fiber stimulation during reaches (Calame et al., 2023), and fits into the framework that consistent and predictable consequences of behavior are learned and compensated by the cerebellar circuitry (Sawtell, 2017). Further work is required to decipher the mechanism of the compensation reported here, and which is typically associated with plasticity at the parallel fiber - PC synapse (Sawtell, 2017).

Variations in swing length dynamically impose modulations of the firing rate transients reported here. These changes in swing-stance transition triggered MLI activity are most likely inherited from changes in granule cell activation and mediated by parallel fibers, as the sensorimotor context of a task is thought to be encoded by granule cell activity (Albus, 1971; Marr, 1969). Depending on the relative weight of direct excitatory inputs to PCs from parallel fibers and inhibitory input through the parallel fiber - MLI pathway to PCs, MLI-mediated inhibition can allow to independently regulate both the baseline activity and the magnitude and direction of firing rate modulation in Purkinje cells during swing-stance transitions. Whether the change in the fraction of modulated cells across learning (sessions) is due to plasticity at the parallel fiber – MLI synapse (Mapelli et al., 2015) remains to be determined.

While classic models have focused on the granule cell to Purkinje cell pathway as the key site of sensorimotor prediction (Sawtell, 2017), the MLI encoding of limb events across locomotor learning could point to a more active role for cerebellar interneurons. This possibility is further supported by the fact that MLI firing rate modulations occured most frequently at stance onset when the mouse terminates a reaching movement for a chosen rung, as recent work has shown that the cerebellar nuclear neurons targeted by Purkinje cells in the simplex bidirectionally encode endpoint velocity in a reaching task (Becker and Person, 2019). Future explorations including recordings of cerebellar input - from mossy fiber, granule cells - and output signals - from Purkinje cells - of the cerebellar cortex during the LocoReach task will further clarify how cerebellar circuit processing enables precise motor coordination. An open question is whether the observed encoding patterns are consistent across MLI subtypes, or differ between stellate and basket cells or MLI2 (Lackey et al., 2023). Our loose cell-attached recordings may have over-sampled stellate cells located superficially. Future work could examine encoding specificity of MLIs leveraging genetic access to subclasses of MLIs.

Locomotion in our task was driven by the treadmill motorization rather than self-initiated. Although this setup constrains forward velocity, the requirements for accurately reaching to the next rung and generating appropriately timed swing–stance transitions remain comparable to free locomotion on runged surfaces. Thus, the core sensorimotor computations underlying paw placement are expected to be largely preserved across both conditions. At the same time, we acknowledge that MLI encoding could differ between treadmill-driven and spontaneous, voluntary, or goal-directed locomotion, particularly given cerebellar contributions to processing reafferent signals. These potential differences warrant future investigation.

The newly developed LocoReach task allows for the simultaneous study of high precision motor output and neural activity. Here, we focus on recording spiking activity of single MLIs. The locomotor task is continuous but we find that MLI activity is linked to specific instances in the stepping cycle, *i.e.*, situations with heightened demands on the precision of locomotor output. Such intermittent engagement across the population has not been reported before and proposes a new role of MLIs in locomotion. This discovery promises new exciting avenues for understanding cerebellar and cerebral function during complex locomotion using LocoReach.

## Methods

Experimental procedures complied with animal care guidelines of the Université Paris Cité. Experimental procedures were approved by the French Ministry of Higher Education, Research and Innovation (APAFIS#11158-2017041316199 v8) in agreement with the European Directive 86/609/EEC and by the ethical committee of the Université Paris Cité.

### LocoReach setup

The LocoReach setup allows the study of the fine details of performance, motor coordination, and learning during skilled locomotion on a treadmill with rungs. The custom treadmill comprised 48 rungs (3 mm diameter, 80 mm long) equally spaced every 1.65 cm (Fig. 1A; outer circumference 80.4 cm) and was 3D printed in semi-transparent material (WaterShed XC 11122) through Stereolithography (Protolabs Euro Services Limited). The half-open treadmill design allowed to place a custom laser-cut acrylic mirror at a ∼ 45*^◦^* angle in the wheel, under the animal. The wheel rotated freely and allowed for self-paced locomotion. For motorization, a servo motor (Futaba S3004, servo 1 in Fig. 1A) moved into a gear and another continuous servo motor (Parallax Continuous Rotation Servo, #900-00008; servo 2 in Fig. 1A) imposed a constant baseline speed of 7.49 cm/s, and 5.8 cm/s in the musciol/saline experiments. On flat surfaces for comparison, mice exhibit the slowest gait pattern ‘walk’ - *i.e.* at least three paws simultaneously on the ground - with average speeds of 9 ± 5 cm/s (± SD) but can reach maximal moving speeds of up to 100 cm/s (Bellardita and Kiehn, 2015; Machado et al., 2015). The rotation of the wheel was recorded with a rotary encoder (CUI Devices, AMT112S-V, resolution: 2048 pulses per rotation), digitized at 40 kHz with a standard DAQ acquisition board (National Instruments, PCIe 6323) and acquired with ACQ4 (Campagnola et al., 2014). Mice were illuminated with infra-red light (a ring of 10 LEDs of 940 nm - Thorlabs LED940E - located around the objective of the camera), videographed at 200 frames/s (camera: Dalsa Genie Nano, NANO-M800-NIR; objective: Fujinon DF6HA-1B; positioned 22 cm from the center of the treadmill) and videos were acquired with ACQ4. An array of four flashing infra-red LEDs (Thorlabs LED940E) was placed to appear in the upper right corner of the behavioral videos (Fig. 1B) and used to synchronize video frames with the electrophysiological- and rotary encoder recordings. Transparent acrylic side walls restricted the animal’s position to the center of the runged surface of the treadmill (Fig. 1A).

### Animals

For the electrophysiological recordings, we used transgenic animals (five males and six females, aged 1-3 months) issued from a cross of mice carrying the GCaMP6f transgene in the Igs7 locus (Ai93(TITL-GCaMP6f)-D; Jackson laboratory stock #024107) and B6 PV-Cre mice (B6.129P2-*Pvalb*^tm1(cre)Arbr^/J; Jackson laboratory stock #017320). B6 PV-Cre mice were used in the optogenetic manipulation experiments (8 animals in total, 3 females and 5 males). All animals lived under inverted day/night cycle with ad libitum access to food and water in enriched environments.

### Chronic window with access port and head-plate surgery

Mice were prepared for electrophysiological recordings during locomotion by implanting a chronic window with silicone access port (Roome and Kuhn, 2014) and fixing a head post. Before surgery (10–15 min), mice were injected with buprenorphine (0.35 mg/kg intraperitoneal) for pain reduction. Lidocaine was used as a local anesthetic before skin opening. Surgeries were performed under isoflurane (1.5–2.0 % by volume in air) to maintain a surgical plane of anesthesia as determined by non-responsiveness to toe pinch. Eye ointment was applied under anesthesia at the start of surgery. Animals were held in a stereotaxic apparatus using ear and nose bars to stabilize the head. Body temperature was set with a heating pad and controller at 37 °C using biofeedback from a rectal thermocouple.

For the surgical procedure, the area of the head over the intermediate cerebellum was first shaved and the underlying skin was treated repeatedly with ethanol (70 %) and Vetedine solution (Centravet) in alternation. The skin above the skull was removed and the opening of the periosteum as well as any membranous tissue were cleaned. Once the skull was dry, a cranial window was carefully drilled using a handheld high-speed micromotor drill (Foredom, FST burr #19007 − 05 with 0.5 mm tip diameter). The center of the craniotomoy was located −6 mm AP and 2 mm ML targeting the left lobulus simplex. Skull removal was performed under application of an ACSF solution to minimize the risk of damaging the dura overlying the brain. After skull removal, the ACSF solution was used to clean any debris from the brain surface and to maintain the brain moist providing an interface to the chronic window. A chronic window was fabricated from a cover glass (#1, thickness 150 *µ*m) laser cut to a small oval shape (3 mm x 2.6 mm) to better fit the cranial opening, and to feature a central hole of ∼ 500 *µ*m diameter for electrode entry. Prior to surgery, the cut hole was sealed with Silgard 184 (Roome and Kuhn, 2014). The window was placed over the cerebellar lobulus simplex and kept in direct contact with the brain by lowering the window with the help of a needle mounted in the stereotaxic setup. The chronic window was fixed in place with dental cement (Super-Bond C & B). After setting of the dental cement, the bone surrounding the craniotomy was thinned with the micromotor drill to allow for better access with the 20x objective and recording electrode. Then a custom-made stainless steel head post with a central opening for the craniotomy was cemented in place with dental cement (Super-Bond C & B). The chronic window was covered with silicone for protective purposes after surgery. Mice were carefully monitored during surgical recovery, with all animals showing normal behavior including the absence of motor deficits. Mice were allowed to recover at least 2 days before proceeding to the locomotion and electrophysiological recording experiments with ad libitum access to food and water.

### LocoReach learning protocol

Prior to locomotor sessions, mice were habituated to the LocoReach setup and the head-fixation (attachment of the surgically implanted head-post to a rigid fixture) on top of the treadmill. For habituation, the treadmill’s surface was covered with a plastic mesh (square grid of 5 × 5 mm) allowing for easy grip. Head-fixation time was gradually increased during four habituation sessions spread over two days, and wheel motorization (same as described below during locomotor sessions) during head-fixation was introduced at the third habituation session.

Mice were positioned with the tip of the nose ∼ 5 mm above the spherical treadmill, and with an angle of ∼ 10*^◦^* between the tangent of the treadmill and the bottom of the lower jaw, a position which was inferred from inspecting free walking on the treadmill. One session consisted of multiple (typically 5, range: 1 − 8) 1 min recording trials which were repeated every 2 – 4 min, *i.e.*, with rest periods of at least 1 min between trials. The speed and the duration of locomotion during the recordings were imposed in an attempt to minimize variations across recordings. The forced locomotion scheme was as follows: recordings started with a 6 s long period of baseline during which the mouse was able to freely walk on the treadmill. At 6 s, servo motor 1 moved the continuous rotation servo 2 in a gear attached to the axis of the treadmill (Fig. 1A). The continuous servo started to linearly accelerates (2.5 cm/s^2^) the wheel at 7 s until the constant baseline speed of 7.49 cm/s was reached at 10 s. This rotation speed was kept constant for 42 s until 52 s at which point the motor linearly decelerated (−2.5 cm/s^2^) until standstill at 55 s and then moved out the gear from the treadmill axis allowing for self-paced walking.

### Stereotaxic injections, optical fiber implantation and real-time optogenetic manipulation

For the optogenetic interference experiments, B6 PV-Cre mice (Jackson laboratory stock #017320) mice underwent the same surgical procedures as outlined above for the chronic window implantation, up to the point of skull cleaning in the stereotaxic setup. A cranial window of ∼ 500 *µ*m diameter was then carefully drilled using a handheld high-speed micromotor drill (Foredom, FST burr #19007 − 05 with 0.5 mm tip diameter). The center of the craniotomoy was located −6 mm AP and 2 mm ML targeting the left lobulus simplex. Subsequently, B6 PV-Cre mice received unilateral injections in the left lobule simplex of 0.7 *µ*l of pAAV-1-hSyn1-Flex-SIO-stGtACR2-FusionRed-dlox (*N* = 5, 2.0 × 10^12^ genome copies/mL; Addgene: 105677-AAV1), while another group of B6 PV-Cre mice received injections of pAAV-9/2-hSyn1-dlox-tdTomato-dlox-WPRE (*N* = 3, 6.1 × 10^10^ genome copies/ml; VVF Zurich: v663-1). These injections were administered through the craniotomy using a Hamilton syringe (Model 7001, Hamilton) at a rate of 0.1 *µ*l/min, employing a glass capillary needle with a diameter ranging from 14 to 18 *µ*m. The injection site was localized to the left lobulus simplex (coordinates: anterior-posterior −6 mm, medial-lateral 2 mm, dorsoventral 0.35 mm). Following each injection, the needle remained in situ for 5 minutes to mitigate backflow before being slowly retracted. Subsequent to the injection procedure, an optical fiber (core diameter 200 *µ*m, numerical aperture 0.39, Thorlabs Inc.) affixed to a ferrule (1.25 mm diameter) was implanted above the injection site (coordinates: anterior-posterior −6 mm, medial-lateral 2 mm, dorsoventral 0.325 mm) and secured to the skull using dental cement (Super-Bond C & B).

Real-time tracking of the position of the left forepaw was achieved using Bonsai during the behavioral experiments (Lopes et al., 2015). Using the animal video stream, a region of interest was placed to comprise the ensemble of FL positions during the task in Bonsai. Image of this ROI underwent multiple processing steps to extract the center of mass of the paw. Whenever the FL center of mass displacement across consecutive frames exceeded a threshold, which was adjusted to identify swing onset, the laser was triggered for 80 ms through a Bonsai-controlled Arduino. The delay between threshold crossing and trigger signal as well as the duration were adjusted to achieve laser activation during and at the end of the swing phase (Fig. 3D,E).

The behavioral task started four weeks following the virus injection. Before exposure to the LocoReach task, mice underwent the same habituation procedure as outlines above. The optical fiber was connected to the implanted ferrule and the real-time extraction parameters (ROI of the front left paw) were adjusted in Bonsai prior to each locomotor session. Optical stimulation was administered using a laser (405 nm wavelength) coupled to a patch cord fiber (core diameter 500 *µ*m, numerical aperture 0.5, Thorlabs), with an output intensity of 30 mW from the fiber tip.

Verification of viral vector expression through histology

After completion of the behavioral tests, mice were anesthetized with an intraperitoneal injection of ketamine (150 mg/kg) and xylazine (10 mg/kg) and perfused transcardially with a solution of cold phosphate buffered saline (PBS), followed by 4% paraformaldehyde (PFA) diluted in PBS. Brains were removed and post-fixed for 24 hr in 4% PFA. 50 *µ*m thick coronal slices of the cerebellum were prepared using a vibrating slicer (Leica VT 1000S, Leica Microsystems, Germany). After 3 washes in PBS, slices were mounted between slides and cover slip with Vectashield Mounting Medium (Vector lab, Burlingame, CA). Mosaics and stack of images (0.5 *µ*m thickness) were acquired around the lobule simplex using an LSM 710, confocal microscope (Zeiss, Jena, Germany) with a plan-apochromat 20x objective. The confocal pinhole was set to 1 Airy unit. Image stacks spanning 40–50 *µ*m were acquired with a z-step of 1 *µ*m (optical section thickness: 0.5 *µ*m). For quantification of GtACR2-positive cells, we selected the single image plane within each stack containing the highest number of FusionRed-positive somata and quantified positive cells in ImageJ (NIH, Bethesda, MD). Cells were classified visually based on morphology and spatial location: molecular layer interneurons (MLIs) exhibited small somata with thin, multipolar dendrites and lacked a large planar dendritic arbor, whereas Purkinje cells (PCs) were identified by somata located in the Purkinje cell layer and a large soma with a prominent apical dendrite extending into the molecular layer. Viral expression and fiber placement were confirmed for all animals and mapped onto sections adapted from the mouse brain atlas (Franklin and Paxinos, 2008). Illustrative images shown are single-plane images selected from the corresponding stacks to reflect the plane with the highest number of GtACR2-positive cells.

### Cell-attached recordings

We conducted cell-attached recordings of the electrical activity of MLIs and PCs in the left lobulus simplex of PV-cre/Ai93(TITL-GCaMP6f)-D mice during the acquisition of the LocoReach task. These mice expressed GCaMP6 in ∼ 50 % of MLIs throughout the cerebellar cortex. We used long-shaft patch pipettes (5 − 10 MΩ resistance) loaded with Alexa 594 (50 *µ*M) in ACSF (150 mM NaCl, 2.5 mM KCl, 1.5 mM CaCl_2_, 1 mM MgCl_2_, 10 mM Hepes, pH 7.3, 280 − 300 mOsm), and performed two-photon microscopy-guided recordings of MLIs identified through their molecular layer localization and by GCaMP6 expression through a chronic window with a silicone access port that allowed for repeated electrode penetration over days (Figure 4A,B; details of the custom build 2-photon laser scanning setup can be found in Franconville et al., 2011). During the recordings, a ground electrode was inserted in an ACSF bath on top of the chronic window. We performed visually guided electrode approaches in about one third of the MLI recordings (22 visually guided recordings out of the 64 MLIs recorded in total; green points in Fig. A.4A). Whenever optical access was compromised due to imaging depth or obstruction (*e.g.* through a blood vessel), we performed blind cell-approaches and recordings. These recordings required to identified cell identity through clustering based on the cell’s electrophysiological signature as MLIs or PCs (Fig. Figure A.4, see below for details).

The current traces of the cell-attached recordings were recorded in voltage-clamp mode using a Axopatch 200B amplifier (Molecular Devices), digitized at 40 kHz (National Instruments, PCIe 6323), and acquired with ACQ4 (Campagnola et al., 2014). Raw electrophysiological traces were analyzed in P-sort to detect spikes of MLIs and simple as well as complex spikes for Purkinje cell recordings (Sedaghat-Nejad et al., 2021). Current traces were band-pass filtered (0.5 to 5 kHz) and spikes clustered using the build-in nonlinear dimensionality reduction in P-sort.

For validation of the optogenetic inactivation, three B6 PV-Cre mice were injected with pAAV-1-hSyn1-Flex-SIO-stGtACR2-FusionRed-dlox into the left lobule simplex two weeks before headplate implantation and chronic window surgery. After three days of recovery, two-photon-guided cell-attached recordings were performed from MLIs expressing GtACR2-FusionRed, with an optic fiber placed in the ACSF bath above the chronic window. Once a cell with stable spontaneous activity was identified and the mouse was stationary, light pulses were delivered using different stimulation protocols; for the example shown in Fig. 3C, we applied 5 Hz laser pulses of 100 ms duration with 200 ms inter-pulse intervals, at 73 mW power, repeated 20 times.

### Clustering of all recorded cells into MLIs and PCs using t-SNE

To further distinguish between MLIs and PCs, we used t-distributed stochastic neighbor embedding clustering based on the average action potential waveform (interval [−1.2, 2] ms, sampling interval 0.05 ms), the spike auto-correlogram (interval [−20; 19] ms, bin size 1 ms), the firing rate, the trough-to-peak delay of the average spike waveform, the spike-count coefficient of variation for multiple bin sizes (0.05 s, 0.5 s, 1 s and 5 s), and the existence of complex spikes as binary variable (Fig. A.4). All these electrophysiological measures where gathered in a design matrix where each row would correspond to a recording and the row would contain the stacked electrophysiological data mentioned in an one-dimensional array. This design matrix is then used in the unsupervised t-SNE method with the following parameters : n_components= 2, perplexity= 45, early_exaggeration= 18. This method identified two separate clusters containing putative MLIs and PCs, respectively. We required the clustering to correctly classify all visually identified cells as MLIs and used this requirement as a criterion to assess clustering performance and fine-tune the clustering parameters. The MLI cluster contains 64 cells from a total of 11 mice, most with 5 trials per cell per recording session (Fig. A.4A inset). The PC contains 34 cells from 11 mice. We discontinued recordings for a given mouse when it reached 10 recording sessions or when the chronic window quality decreased due to dura regrowth or tissue thickening. Cells with unstable spike signals (*e.g.*, complete disappearance of spikes for prolonged periods) during the 60 second trial period were excluded from the analysis.

### Paw tracking and swing-stance segementation

The videos recorded during the locomotor sesions were analyzed using DeepLabCut (DLC; Mathis et al., 2018) to extract the x- and y-positions of front left (FL), front right (FR), hind left (HL) and hind right (HR) paws across individual frames in the bottom view of the animal (Fig. 1B,C). Multiple DLC relabelling and retraining iterations were performed to achieve a target error rate of *<* 0.0005 % per video, where errors were identified when paw displacement speed across successive frames exceeded *>* 60 pixels/s. Individual frames which exceeded this threshold were excluded from further analysis. Using the congruent paw- and wheel speed periods during stance and the known wheel speed at the surface of the treadmill, we transformed the paw speed extracted from the videos into cm/s, using the scaling factor 0.025 cm/pixel. A custom-written Python script was used to track the position of all rungs present in each video frame, based on the fixed and consistent spacing between the rungs. The relative difference between paw x-speed and wheel speed (speed difference *>* 10 cm/s for at least three consecutive frames), as well as the paw-to-rung distance (paw and mouse specific based on the histogram of paw-rung distances) were used to segregate the paw trajectories into swing and stance phases (Fig. 1D). From these segmented traces, we quantified swing (or stride) length and duration as the spatial distance and temporal interval, respectively, between the stance-to-swing transition and the subsequent swing-to-stance transition. Miss steps were extracted based on multimodal paw speed profiles during the swing face indicative of alternating deceleration and acceleration phases due to attempted rung approaches (Fig. 8A). That is, if the relative paw-speed (paw speed - wheel speed) dropped below 10 cm/s for at least three consecutive values (excluding the beginning and the end of the swing) or for at least three times, the swing was classified as a ‘miss step’. The swing–stance segmentation and miss step identification parameter values were determined through visual inspection of the trajectories and step classifications on a large subset of steps in the video recordings. Using extracted paw positions, paw speed, swing-stance phases, rung locations and miss steps, we computed the stride- and inter-paw coordination parameters shown in Fig. 2, 3 and A.1. All behavioral quantification focuses exclusively on the constant-speed period between 10 and 52 s, excluding the acceleration and deceleration phases.

### PSTH analysis

Peristimulus time histograms were computed based on consecutive bins (bins size 20 ms) of spike times in the interval [−300, 400] ms centered on all swing- or stance onsets of a recording, and normalized by the number of events and the bin size to yield average firing rate. Shuffled PSTHs are obtained through jittering the original spike train by adding to each spike an independent Gaussian random variable of zero mean and *J* = 500 ms standrard deviation (STD), which eliminates correlations to behavior on time-scales ≪ *J*. The mean shuffled PSTH, the 5th and 95th percentiles and the STD along the interval [−300, 400] ms was computed from 300 shuffled PSTHs obtained with different random number seeds. Individual recordings were classified as significantly up/down modulated before or after an event, if the PSTH crossed the 95th/5th percentile of the shuffled PSTHs for at least two consecutive values in the 100 ms interval preceding or following stance- or swing onset ([−100, 100] ms window). The z-scored PSTH was calculated using the shuffled PSTHs according to 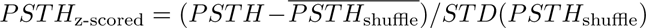. And the area under the curve (AUC) of the z-scored PSTH corresponds the integral of absolute values over the interval [−100, 0] ms for before, [0, 100] ms for after stance- or swing onset, and over [−100, 100] ms for the combined AUC.

For the swing-length and swing-duration percentile-based analysis (Figs. 6, A.8, A.9), all swings of a given cell during one session of multiple trials where subdivided according to the percentile ranges [0 − 20], [20 − 40], [40 − 60], [60 − 80], [80 − 100] of all swing lengths or durations. The z-scored PSTH was then calculated for swings falling withing a specific percentile range. The AUC and whether a cell in a given percentile range was modulated or not was evaluated as specified above.

### Event kernels and linear models

To measure the influence of swing- and stance-onsets on MLI firing rates, we fitted L2- and L1-penalized linear models in Python, implemented in the package scikit-learn ("Ridge" and "Lasso", respectively). Firing rates were calculated from binned counts with a bin size of 10 ms. We detrended firing rates by subtracting smoothed signals (Gaussian filter with *σ* = 300*ms*) and subsequently smoothed rates using a Gaussian kernel (*σ* = 20*ms*). After selecting motorized periods, we z-scored and concatenated all recordings from the same cell.

Multistep linear regression was performed by first constructing the design matrix for event kernel estimation: For each limb and event type (swing vs. stance onset), we included a set of 31 time-shifted delta-function regressors that spanned the peri-event period from 150 ms before to 150 ms after each event, resulting in a 248(31 × 2 × 4) × *N_timepoints_* design matrix. Event kernels were obtained as the set of 31 L2-regularized weights for each limb and event.

For the sparsity-constrained second regression step, we normalized event kernels to have maximum absolute weights equal to 1. We then convolved kernels with the indicator function for the respective event, resulting in a 8 × *N_timepoints_* design matrix for 4 limbs and 2 event types. For analyses including interaction terms (event × trial number, event × swing length, event × swing duration), we multiplied all 8 event regressors with the associated value, resulting in a 32 × *N_timepoints_* design matrix. In Fig. A.12, we also included 8 continuous regressors (x-positions for each limb, and wheel-velocity corrected speed along x for each limb). We then estimated L1-regularized weights based on the new design matrix.

Cross-validation was performed to determine optimal L2- and L1-regularization hyperparameters. We first chunked time series into pieces of 90 ms. To decorrelate adjacent data points (which are correlated due to Gaussian smoothing), we removed the first and last 20 ms from each chunk of data. We then randomly assigned chunks to train and test set (80% vs. 20%). Optimal L1- and L2-parameters were determined based on the maximal model *R*^2^ on test data. Cross-validated *R*^2^ was also used to determine the significance of single regressors: For each event, we shuffled temporal chunks of 200 ms of the regressor of interest before performing the L2-regularized regression step. For interaction terms, we shuffled the respective modulating variable (trial number, swing length, swing duration) after performing L2-, but before performing L1-regularized regression. To measure the unique contribution of a variable (Δ*R*^2^), we calculated the difference in *R*^2^ between the model where the variable of interest was shuffled vs. the *R*^2^ on the full (unshuffled) model. Single-variable *R*^2^ values were measured after shuffling all variables but the variable of interest (Musall et al., 2019). We repeated cross-validation 100 times on random train/test splits to calculate 95% confidence intervals for each measure. Variables were considered to have a significant effect if confidence intervals of single-variable *R*^2^ or Δ*R*^2^ were above zero (Figs. 7, A.11 and A.12).

To compare model performance between discrete event-based basis-function regression models and continuous regression models, we fitted (via the above-described two-step L1-/L2-procedure) models containing 8 event kernels, to which we added 8 continuous regressors, corresponding to x-positions and x-velocities of each limb. Collinearity between continuous and discrete regressors was quantified as the Pearson correlation coefficient of L2-regularized event kernel regressors and the continuous variables (Fig. A.12A). Cross-validated *R*^2^ values were obtained for the full model, for a model with shuffled continuous regressors, and for a model with shuffled discrete event-based regressors (Fig. A.12B). For 8D-F, we fitted models containing only discrete or only continuous regressors, respectively. We aligned MLI neural activity to actual and expected stance onset around correct and missteps, and calculated average firing rates in the respective peri-event window. In comparison, we show average predicted MLI activity from the event-based regression model and the continuous regression model. We quantified fits around the respective events by measuring the explained variance as the non-cross-validated *R*^2^ from 150 ms before up to true/expected stance onset, respectively (Fig. 8E,F). The choice of comparing non-cross-validated fits is justified by the infrequent occurrence of missteps, making cross-validation specifically around single-paw missteps technically infeasible. Panel E and F in Fig. 8 compare the relative drop in the *R*^2^ from correct to mis-steps in the discrete and the continuous regressor model. Since these values are not cross-validated and a better performance of the basis-function model is expected due to the higher number of parameters, only the interaction effect between model and event type should be interpreted.

Early vs. late learning categorization was performed solely on the basis of sessions: For each mouse with *N* sessions, the first *N/*2 sessions were labeled as “early learning”, and the latter *N/*2 sessions were labeled as “late learning”; for odd numbers of sessions, we included the middle session in “early learning”. After this categorization, we excluded cells with a negative cross-validated *R*^2^. This criterion led to an unequal number of cells for early vs. late sessions.

### Statistics

Statistical differences in locomotor parameters were analyzed using mixed linear models with session, trial, paw as fixed effects variables and animal as a random effect (*parameter* = *session* + *trial* + *paw* + (1|*animal*)). Normality of the residuals was verified with D’Agostino’s test for mixed linear models. To test for linear relationship between variables, we used Pearson’s correlation or linear least-squares regression. To test for differences within a group, between groups or if a group mean is different from zero, we used paired, unpaired or one sample Student’s t-test respectively. Normality was tested with D’Agostino’s test, homogeneity of variance tested with Levene’s test and One-way ANOVA was performed to test the changes in the PSTH AUC for swing length percentiles. Results were considered statistically significant when *p <* 0.05. The main effect and statistical significance are given in the appropriate figure legend. These statistical tests were implemented in custom Python scripts (Python version 3.8) using the Scipy (version 1.8.1) and the Statsmodels (version 0.13.2) modules.

## Data availability

The datasets generated during and/or analysed during the current study are available from the corresponding author on reasonable request.

## Code availability

All custom written Python scripts used to analyze the data and generate the manuscript figures are available from the corresponding author on reasonable request.

## Acknowledgements

We thank Dr. Isabel Llano for careful reading and feedback on the manuscript. We thank Drs. Kishore Kuchibhotla, Florent Haiss and David DiGregorio for their insightful comments on the manuscript. This study was supported by the ‘Agence National de la Recherche’ (ANR, project : “WalkingCrossingNeurons”, ANR-18-CE37-0006-01; and project: “FrontCog”, ANR-17-EURE-0017), and an EMBO postdoctoral fellowship to Heike Stein (ALTF 471-2021). We acknowledge the use of the animal facilities and the prototyping platform at BioMedTech Facilities (https://biomedicale.u-paris.fr/biomedtech-facilities/, INSERM US 36, CNRS UAR 2009, Université Paris Cité). We thank Patrice Jegouzo and Thierry Bastien of the prototyping platform at BioMedTech Facilities for manufacturing parts of the LocoReach setup. We thank Sandrine El Marhomy (Université Paris Cité, CNRS, SPPIN, Paris, France) for managing the mice colony and for genotyping.

## Author Contributions Statement

M.G. conceived and designed the study. M.G. and J.G. built the setup and performed first pilot experiments. A.A. carried out the experiments presented here. A.J. conducted the post mortem histology. A.A., C.B. and M.G. carried out the analysis. H.S. and N.A.C.G. contributed the GLM analysis. M.G., A.A., H.S. and N.A.C.G. discussed results, interpreted the data and wrote the manuscript. M.G. supervised the project.

## Competing Interests Statement

The authors declare no competing interests.

## Supplementary Information

**Supplementary Video 1. Video illustrating the contribution of somatosensory inputs during LocoReach task execution.** Playback is slowed down 10×. Whisker movements were recorded at 250 fps. whiskerVideo.mkv (video properties are codec 4CC: FFV1; image size: 1920×1080; frame rate: 25 fps; average bitrate: 65, 005 kbps; total duration: 1 min 39 s)

### A. Supplementary figures

**Figure A.1:**
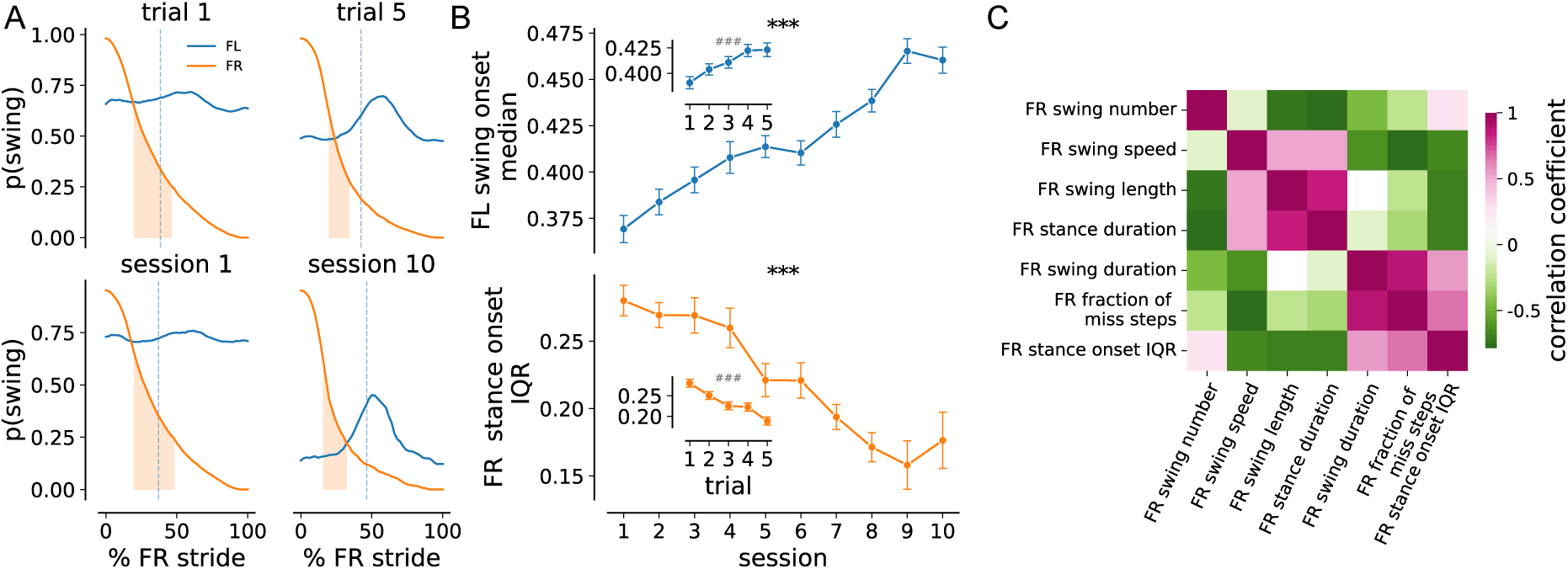
Supplementary: Front paw coordination across trials and sessions with FR paw as reference. (**A**) Average Hildebrand plots aligned to the FR swing onset, showing the probability of swing for FL (blue) and FR (orange) during trial 1 (upper left), trial 5 (upper right), session 1 (lower left), and session 10 (lower right). The orange shaded band indicates the FR stance-onset interquartile range (IQR; 25–75%), and the vertical blue dashed line marks the median FL swing-onset time relative to FR swing onset. (**B**) Coordination metrics referenced to FR swing onset. Top: median FL swing-onset time relative to FR swing onset (session effect: *C* = 0.004, *p <* 0.0001; trial effect: *C* = 0.002, *p* = 0.001). Bottom: FR stance-onset IQR relative to FR swing onset (session effect: *C* = −0.010, *p <* 0.0001; trial effect: *C* = −0.009, *p <* 0.0001). Measures over trials are shown as insets in panels where the trial effect is significant. Averages across animals are shown over sessions and over trials. (**C**) Pearson correlation matrix between behavioral parameters for the FR paw. Colors represent correlation coefficients with *p <* 0.05 (*N* = 11). In (B), data are presented as mean ± SEM. Statistical significance of effects was determined using a linear mixed model. The regression coefficient (*C*) and corresponding *p*-value are reported for each effect tested. *Symbols:* \**p <* 0.05, \*\**p <* 0.01, \*\*\**p <* 0.0001 effect of sessions; #*p <* 0.05, ##*p <* 0.01, ###*p <* 0.0001 trial effect.

**Figure A.2:**
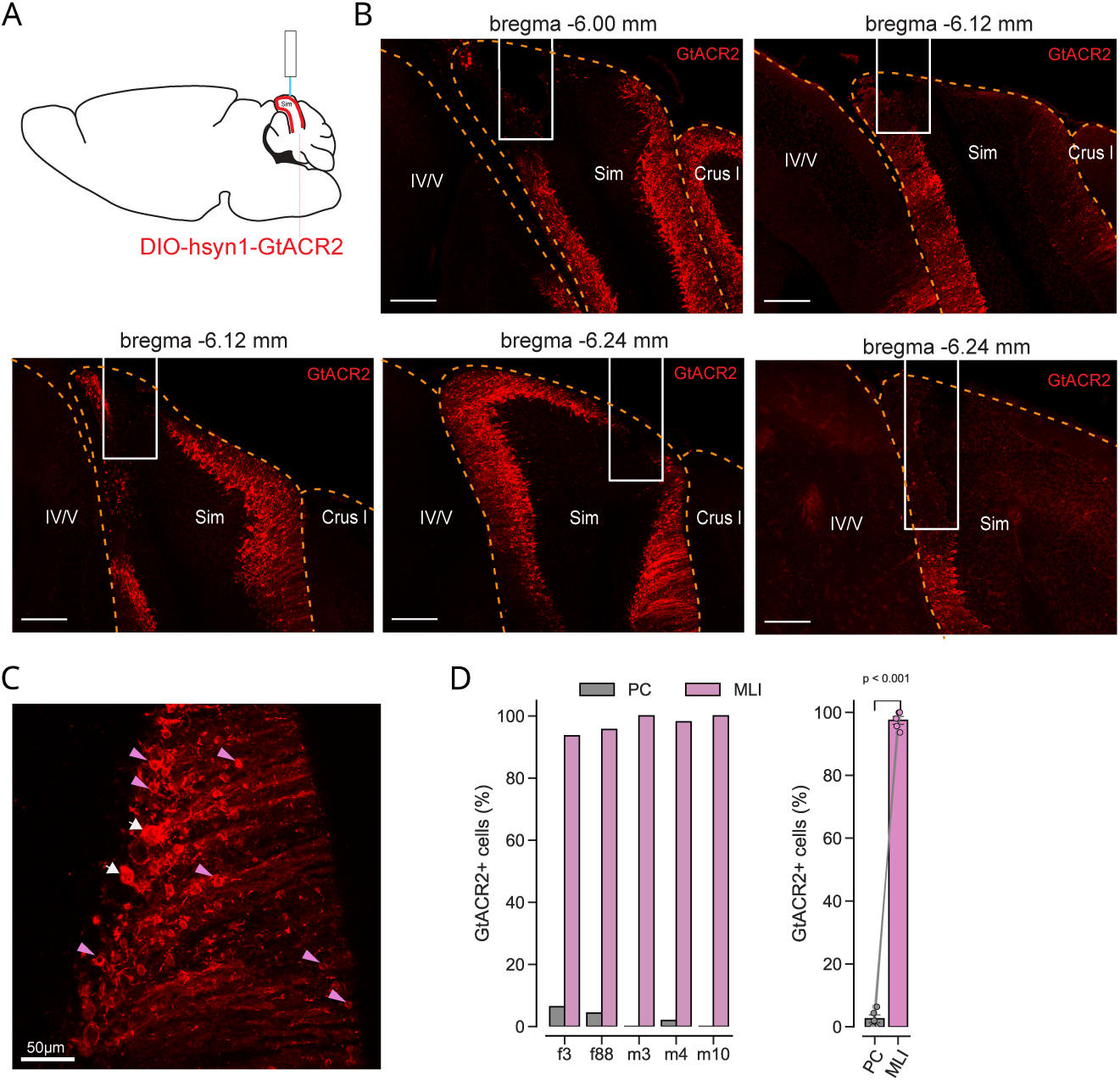
GtACR2-FusionRed expression in all animals of the real-time perturbation experiments. (**A**) Schematic of of the optogenetic perturbation experiments. Blue-light sensitive inhibitory opsin GtACR2 was virally targeted to the lobulus simplex. The opsin was activated during the experiment in a behavioral-dependent manner, *i.e.*, conditioned on the swing phase of the front-left paw. (**B**) Photomicrographs showing spread of GtACR2-FusionRed and implanted optical fiber position in coronal cerebellar sections (scale bar: 200 *µ*m). White squares represent the outlines of the optic fiber implant. Lobulus simplex (Sim), lobule IV/V of the vermis and Crus I are marked. The anterior-posterior position of the shown slice is given on the top of each panel. Each panel represents a different animal subjected to the real-time perturbation experiment (*N* = 5). (**C**) Zoom in photomicrographs showing expression pattern of GtACR2-FusionRed (scale bar: 50 *µ*m). White arrows highlight postitive PCs and pink arrows indicate positive MLIs. Note that not all positive MLIs are highlighted. (**D**) Quantification of PCs/MLIs positive for GTACR2-FusionRed. Manual counting of positive neurons was based on morphology and somata location. Cells were identified as GTACR2-FusionRed positive cells and labeled as MLIs when somata were small (<15 *µ*m), with thin and multipolarized dendrites and lacked a large planar dendrite; cells were identified as PCs when somata were positioned in the Purkinje cell layer and exhibited a large soma with a prominent apical dendrite extending into the molecular layer. Proportion of GTACR2-FusionRed positive PCs (gray) and MLIs (pink) per animal (left panel), the the average across all animals (right panel). Student’s *t*-tests, *t* = 38.026, *p <* 0.001.

**Figure A.3:**
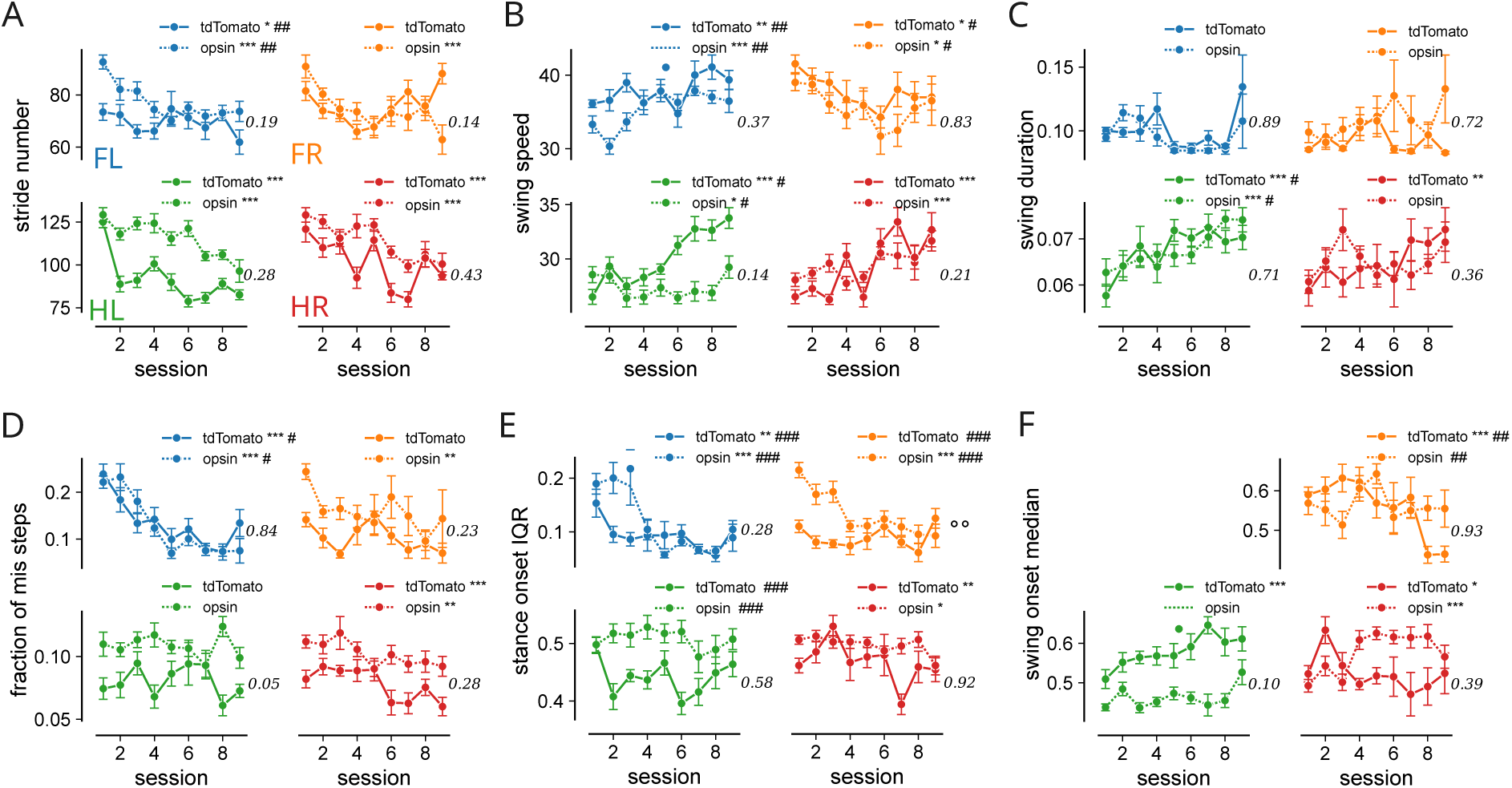
Behavioral measures during real-time inactivation of lobulus simplex. Paw resolved means of behavioral measures are shown across sessions for tdTomato (solid, *N* = 3) and GtACR2 (dashed, *N* = 5) animals. Behavioral measures equivalent to the depiction in Fig. 2 and Fig. 3. (**A**) Swing length. (**B**) Swing speed. (**C**) Swing duration. (**D**) Fraction of miss steps. (**E**) Stance onset IQR. (**F**) Median swing-onset time relative to FL swing onset. Data are mean ± SEM over nine sessions. Statistical significance was assessed with a linear mixed mode and p-values are reported for effects across sessions unless otherwise specified. Symbols: ∗*p <* 0.05, ∗ ∗ *p <* 0.01, ∗ ∗ ∗*p <* 0.0001 (session effects); #*p <* 0.05, ##*p <* 0.01, ###*p <* 0.0001 (trial effects); °*p <* 0.05, °°*p <* 0.01 (treatment effects).

**Figure A.4:**
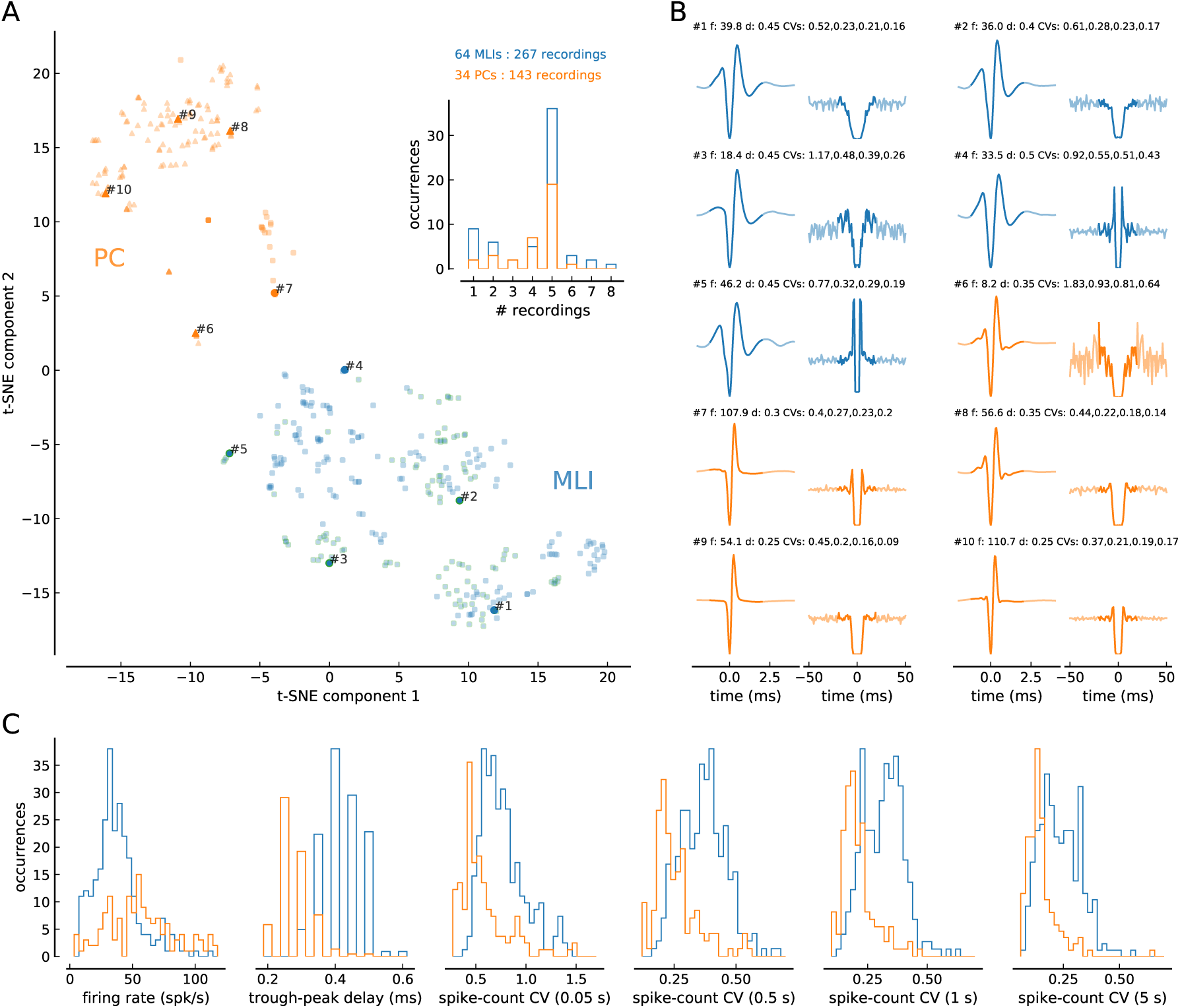
Clustering of all recorded cells with t-distributed Stochastic Neighbor Embedding (t-SNE). (**A**) All recordings are shown based on their first two t-SNE components. t-SNE converts similarities between data points to joint probabilities and tries to minimize the Kullback-Leibler divergence between the joint probabilities of the low-dimensional embedding and the high-dimensional data. For the t-SNE dimensionality reduction, the spike-waveform, the autocorrelogram, the firing rate, the trough-to-peak delay of the waveform, the spike-count coefficient of variation for four different bin sizes (0.05, 0.5, 1 and 5 s), and the existence or not of complex spikes are used as input parameters. Blue symbols are identified as MLIs and orange as PCs. Triangles indicate cells with complex spikes. Symbols with green edge were visually identified during the recording. The inset shows histograms of the number of recordings (trials) performed for all cells in each populations. (**B**) Average spike waveform (left) and autocorrelgogram (right) for ten example cells. The examples are marked in panel (A). Firing rate, *f*, in spk/s and the trough-to-peak delay, *d*, in ms and the spike-count coefficient of variation for four different bin sizes (0.05, 0.5, 1 and 5 s) are shown for each cell (top of the panel). The portion of the curves used for the t-SNE dimensionality reduction is shown in bold color (see Methods for details). (**C**) Histograms of other parameters used in the dimensionality reduction. Each measure is shown for MLIs (blue) and PCs (orange). From left to right : firing rate, trough-to-peak delay of the aveage spike waveform, and the spike-count coefficient of variation for four different bin sizes : 0.05, 0.5, 1 and 5 s.

**Figure A.5:**
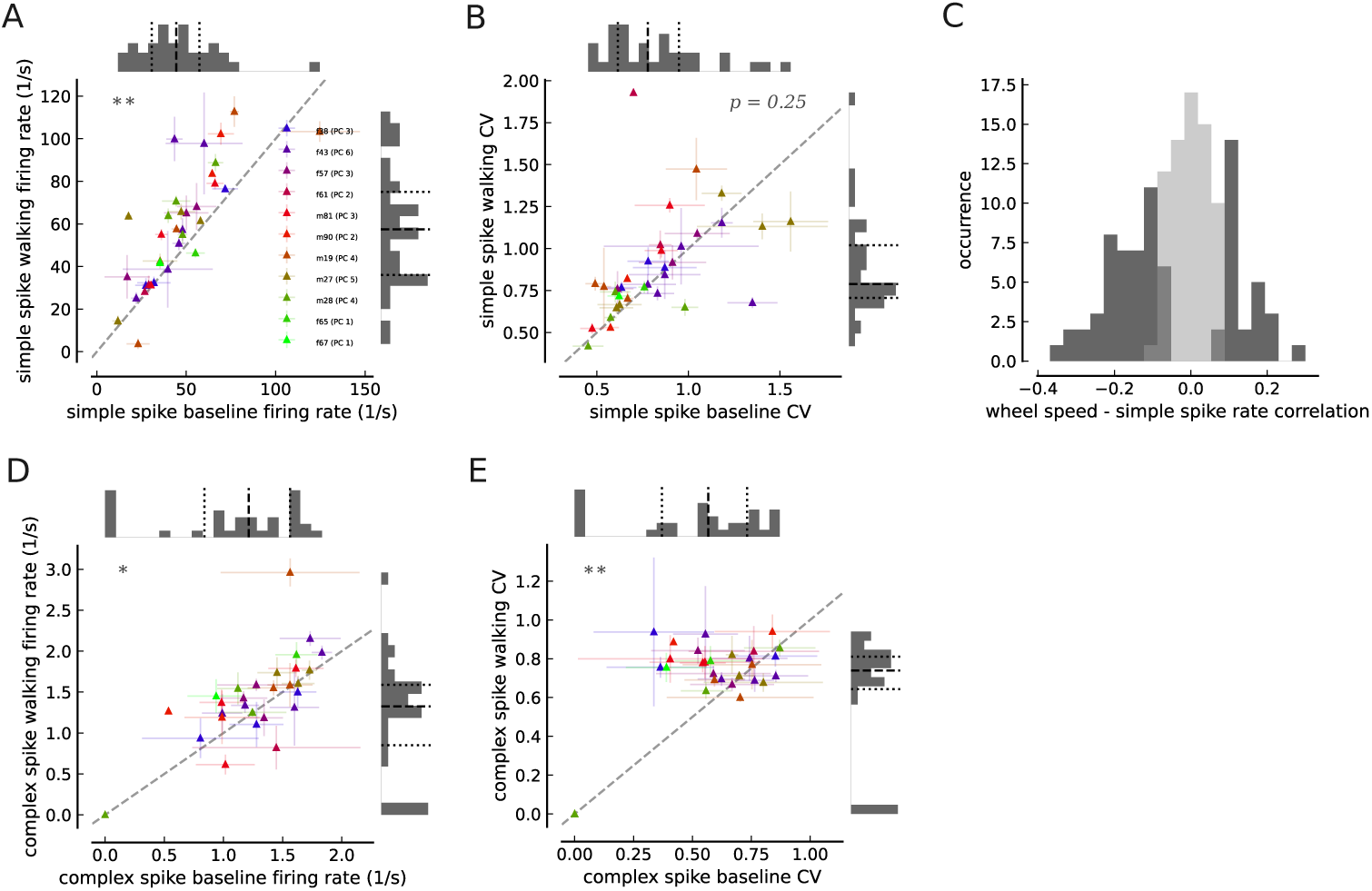
PC activity during LocoReach task. Similar depiction as in Fig. 4H,I but for Purkinje cells simple- and complex spike activity. (**A**) Walking period simple spike firing rate over baseline period firing rate (paired Student’s t-test: *t* = −4.32 *df* = 33, *p <* 0.01). (**B**) Simple spike walking period CV over baseline period CV (paired Student’s t-test: *t* = −1.15, *df* = 97, *p* = 0.25). (**C**) Distribution of walking period correlation between PC simple spike firing rate and wheel speed (*N* = 143 for all recorded trials). Significant correlations (*p <* 0.05) shown in dark gray, and non-significant correlations in light gray. (**D**) Walking period complex spike firing rate over baseline period firing rate (paired Student’s t-test: *t* = −2.25, *df* = 33, *p <* 0.05). (**E**) Complex spike walking period CV over baseline period CV (paired Student’s t-test: *t* = −3.64, *df* = 33, *p <* 0.01). Mean and STD are shown across trials for a given cell. Cells recorded from the same animal in same color (see panel (A) where labels correspond to animal ID and total number of recorded cells per animal in parentheses). Top and right histograms show distribution of respective variable together with 25th (dotted), 50th (dashed) and 75th (dotted) percentiles.

**Figure A.6:**
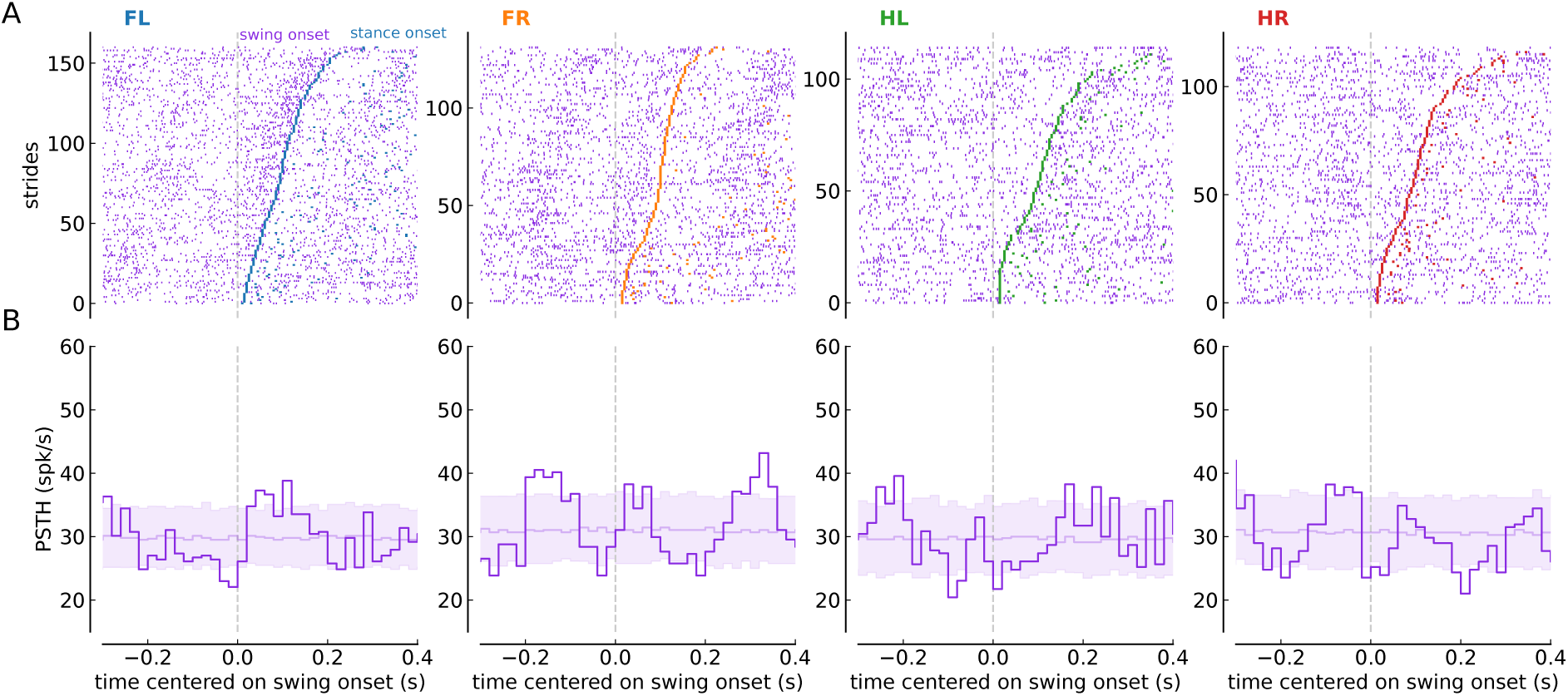
Swing onset-aligned MLI activity modulations. Similar depiction as in Fig. 5B,C but for swing-onset alignment. (**A**) Raster plots showing the occurrence of spikes aligned to swing onset (vertical gray dashed lines) during all strides of each paw (four columns for FL, FR, HL and HR from left), for an example recording of an MLI. Strides are sorted by swing duration. Ordered, colored ticks at positive times mark stance onset time for the swing starting at 0 s, and the following tick marks swing onset for the subsequent swing. (**B**) Swing-onset aligned PSTH corresponding to each paw’s raster plot shown in (A). Pale purple lines and shaded area are mean and [5, 95] percentile intervals of 300 swing-onset aligned PSTHs with shuffled spikes times (see Methods for details).

**Figure A.7:**
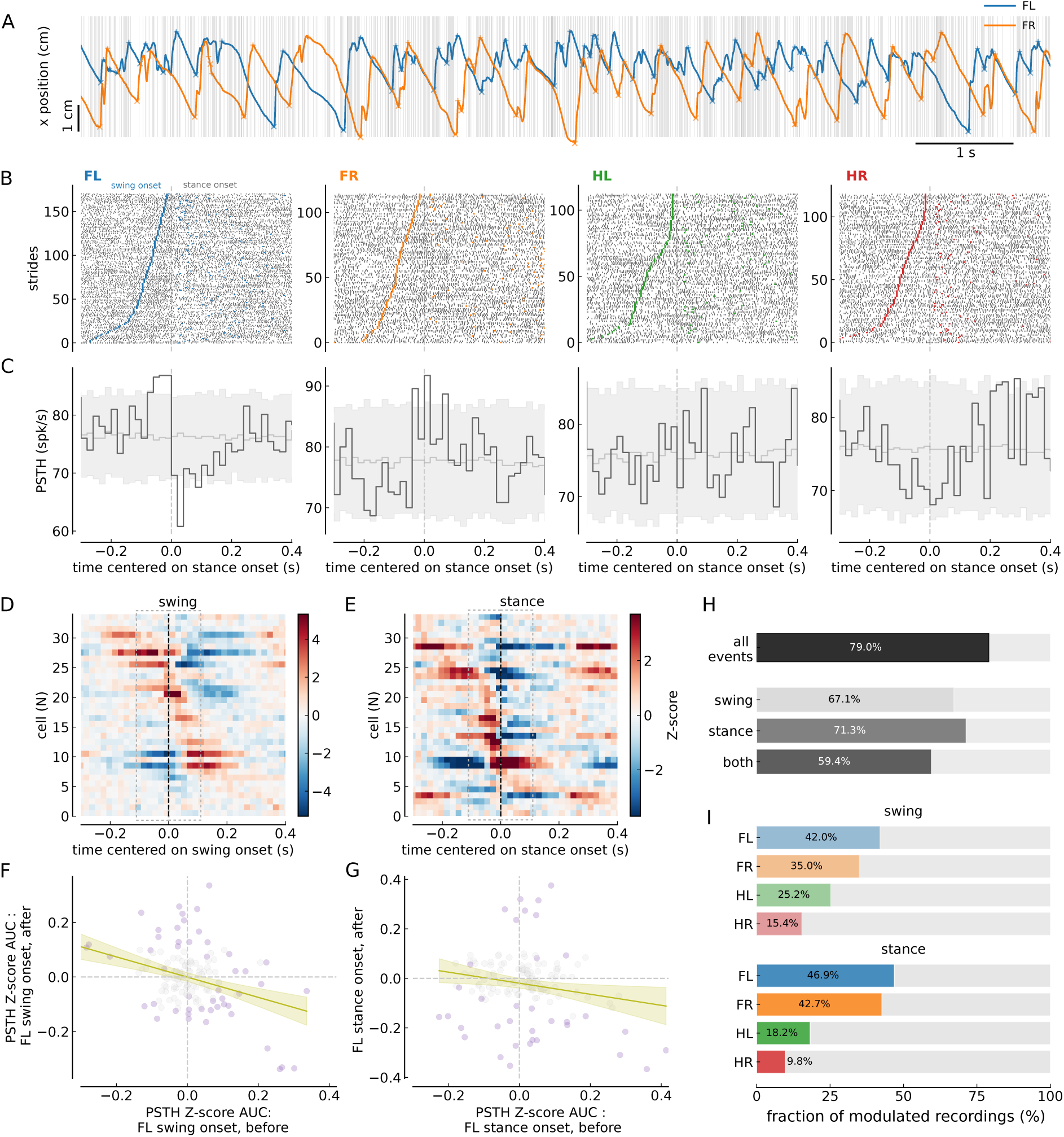
PC simple spike activity modulations swing- and stance-onset aligned. (**A**) Simple spiking activity of an example PC (grey) and simultaneously recorded FL (blue) and FR paw (orange) x-positions during a sample period of a single trial. ‘x’ and ‘+’ symbol represent swing onset and and offset, respectively. (**B**) Raster plots showing the occurrence of simple spikes aligned to stance onset (vertical gray dashed lines) during all strides of each paw (four columns for FL, FR, HL and HR from left), for an example recording of an PC. Strides are sorted by swing duration. Colored ticks at negative times mark swing onset time for the swing ending at 0 s, and at positive times the swing onset for the subsequent swing. (**C**) Stance-onset aligned PSTHs corresponding to each paw’s raster plot shown in (B). Pale gray lines and shaded area are mean and [5, 95] percentile intervals of 300 stance-onset aligned PSTHs with shuffled spikes times (see Methods for details). (**D**) Swing-onset aligned PSTHs of all PC cells (*N* = 34) for the FL paw. PSTHs are shown as z-scores (see Material and Methods) and ordered according to the occurrence of the positive peak, with early peaks on top and peaks occurring later at the bottom. Each PC cell’s PSTH is averaged across all recordings of that cell. (**E**) Same as D but for stance-onset aligned PSTHs. (**F**) The area under the z-scored PSTH curve (AUC) before (x-axis) vs after (y-axis) FL swing-onset. Significantly modulated PSTHs before and/or after are shown with purple circles. The criterion for sig. modulation is two consecutive crossings of the [5, 95] percentile interval of shuffled PSTHs during the 100 ms before or after stance onset. The interval of the AUC evaluation (100 ms) in indicated by gray circles in D and E. A linear regression was performed (Slope = −0.371, Intercept = −0.001, *R*^2^ = 0.153, adjusted *R*^2^ = 0.147, slope p-value *<* 0.001, slope 95% CI: −0.516*…* − 0.226). (**G**) Same as F but for the stance-onset z-scored PSTH curve. Linear regression results : Slope = −0.223, Intercept = −0.019, *R*^2^ = 0.04, adjusted *R*^2^ = 0.033, slope p-value 0.017, slope 95% CI: −0.406*…* − 0.041. (**H**) Bar plots showing the percentage of significantly modulated PCs for all paws and all events (top), for swing- (2nd row), stance-onset (3 row), or both swing- and stance-onset (bottom row). (**I**) Paw resolved fraction of significantly modulated PCs for swing- (top) and stance-onset (bottom).

**Figure A.8:**
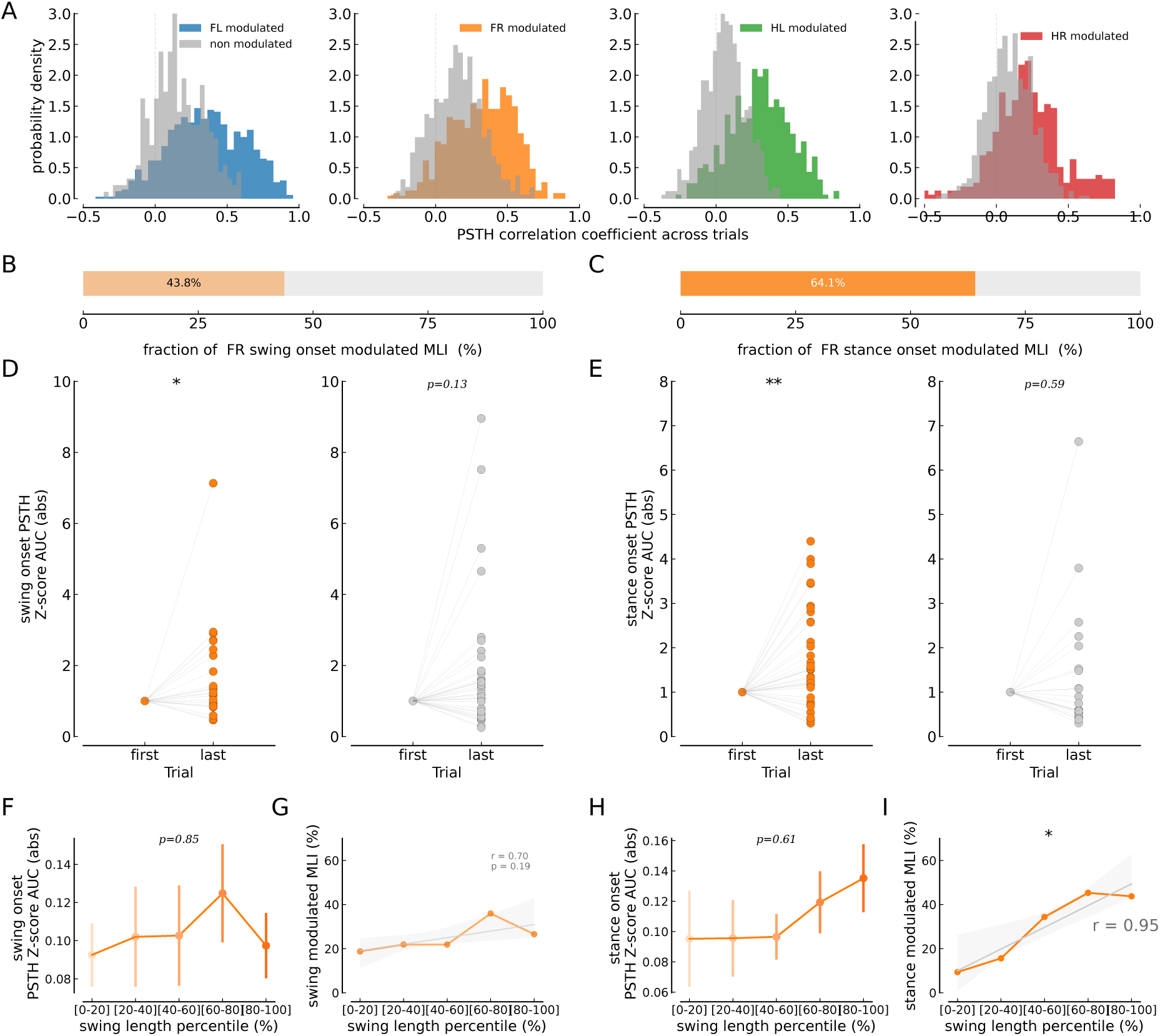
Across-trial FR paw swing- and stance onset-aligned MLI activity. Same depiction as in Fig. 6 but for FR paw (except for (A)). (**A**) Pearson correlation between PSTHs of different trails of a given cell. The distribution of correlations is shown for all four paws (panels from left to right), for modulated cells (in color) and for non-modulated cells (grey). (**B,C**) Fraction of MLIs with at least one or more trials with significant modulations around FR swing (B) or stance (C) onset. (**D,E**) The area under the z-scored PSTH curve (AUC) around FR swing onset (D) and FR stance onset (E) is shown for the last trial relative to the first trial of modulated (left panel) and non-modulated (right panel) cells (analysis window [−100, 100] ms; paired Student’s t-test, *p*-values reported in the panels). (**F,H**) Absolute AUC for swing (F, one-way ANOVA, *F* = 0.336, *p* = 0.852) and stance onset (H, one-way ANOVA, *F* = 0.679, *p* = 0.607) of cells modulated for a given range of swing length percentile. (**G,I**) Fraction of swing onset (G) and stance onset (I) modulated MLIs for ranges of FR paw swing length percentiles (Linear regression analysis, correlation coefficient *r*, and the significance of the regression *p* are reported in the panels). Significance is marked according to \**p <* 0.05 and \*\**p <* 0.01.

**Figure A.9:**
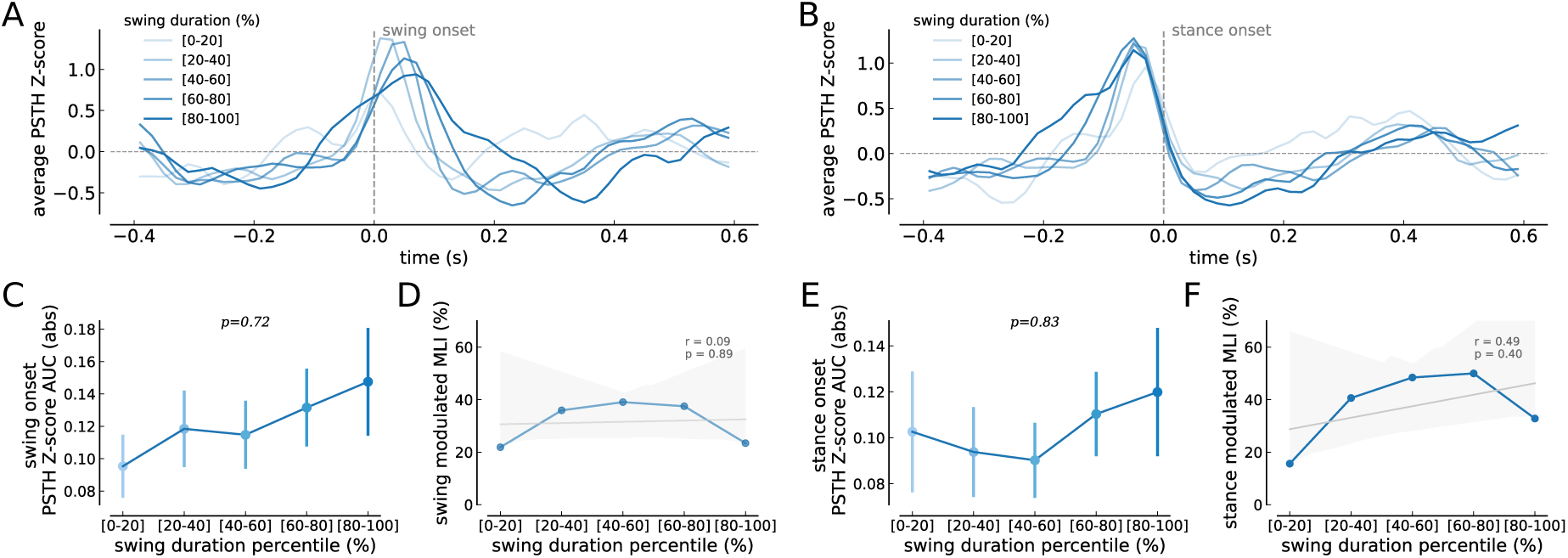
Across-trial FL paw swing- and stance onset-aligned MLI activity for stride durations ranges. Could the change in activity modulation across trials be related to the fact that swings become more stereotyped, since swing duration is less variable later in the session (Fig. 2G,H)? Here, we computed the firing rate modulation strength and the fraction of modulated cells based for swings falling within defined ranges of swing durations during one session (swing duration percentile-based analysis). (**A, B**) Average FL swing (A) and stance (B) onset-aligned, z-scored PSTH of significantly modulated MLIs for increasing ranges of swing length percentiles. (**C,E**) AUC for swing (C, one-way ANOVA, *F* = 0.519, *p* = 0.721) and stance onset (E, one-way ANOVA, *F* = 0.374, *p* = 0.83) of cells modulated for a given range of swing duration percentile. (**D,F**) Fraction of swing onset (D) and stance onset (F) modulated MLIs for ranges of swing duration percentiles (Linear regression analysis, correlation coefficient *r*, and the significance of the regression *p* are reported in the panels). In summary, neither the activity modulation magnitude nor the fraction of modulated cells depended on the swing duration across trials. Interestingly, the peak of the average swing-onset aligned PSTH for modulated cells shifts with the swing length (A), highlighting the fact that the activity modulation is linked to swing end and not the beginning.

**Figure A.10:**
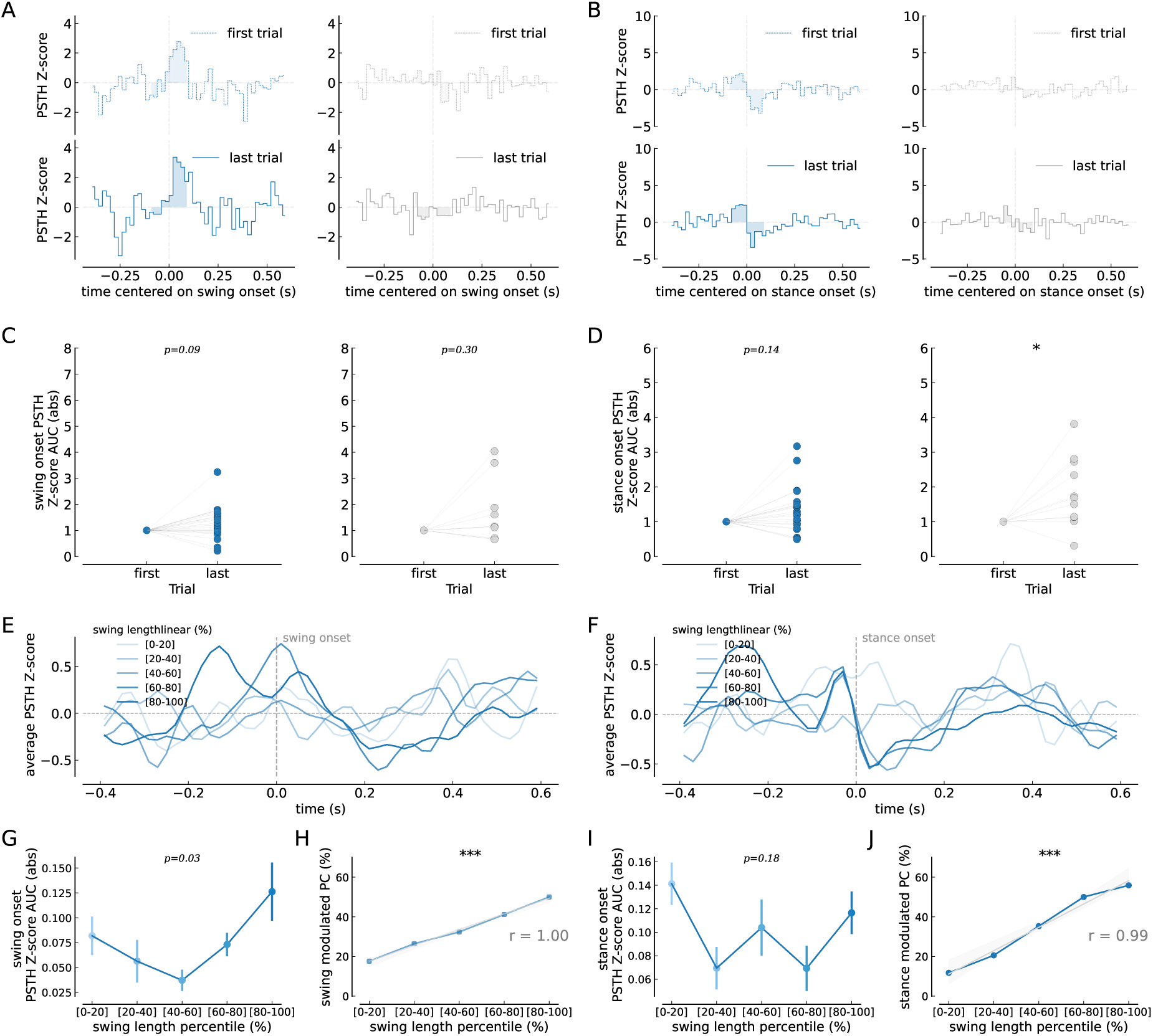
Across-trial FL paw swing- and stance onset-aligned PC simple spike activity correlates with swing length. Same as Fig. 6 but for PC activity. (**A**) Normalized FL swing onset-aligned PSTHs of a significantly modulated (left) and non-modulated (right) example PC around swing onset. The z-scored PSTH for first (dashed line, top panel) and last trials (full line, bottom panel) are shown and the area under curve (AUC) is shown as shaded region (analysis window [−100, 100] ms). (**B**) Same as (A) but for a modulated and non-modulated cell around FL stance onset. (**C,D**) The area under the z-scored PSTH curve (AUC) around FL swing onset (C) and FL stance onset (D) is shown for the last trial relative to the first trial of modulated (left panel) and non-modulated (right panel) cells (analysis window [−100, 100] ms; paired Student’s t-test, *p*-values reported in the panels). (**E**) Average FL stance onset-aligned, z-scored PSTH of significantly modulated PCs for increasing ranges of swing length percentiles. (**F**) Same as (E) but for cells modulated around stance onset (**G,I**) AUC for swing (G, one-way ANOVA, *F* = 2.8190, *p* = 0.034) and stance onset (I, one-way ANOVA, *F* = 1.6458, *p* = 0.176) of cells modulated for a given range of swing length percentiles. (**H,J**) Fraction of swing onset (H) and stance onset (J) modulated PCs for ranges of swing length percentiles (Linear regression analysis, correlation coefficient *r*, and the significance of the regression *p* are reported in the panels). Significance is marked according to \**p <* 0.05, \*\**p <* 0.01 and \*\**p <* 0.001.

**Figure A.11:**
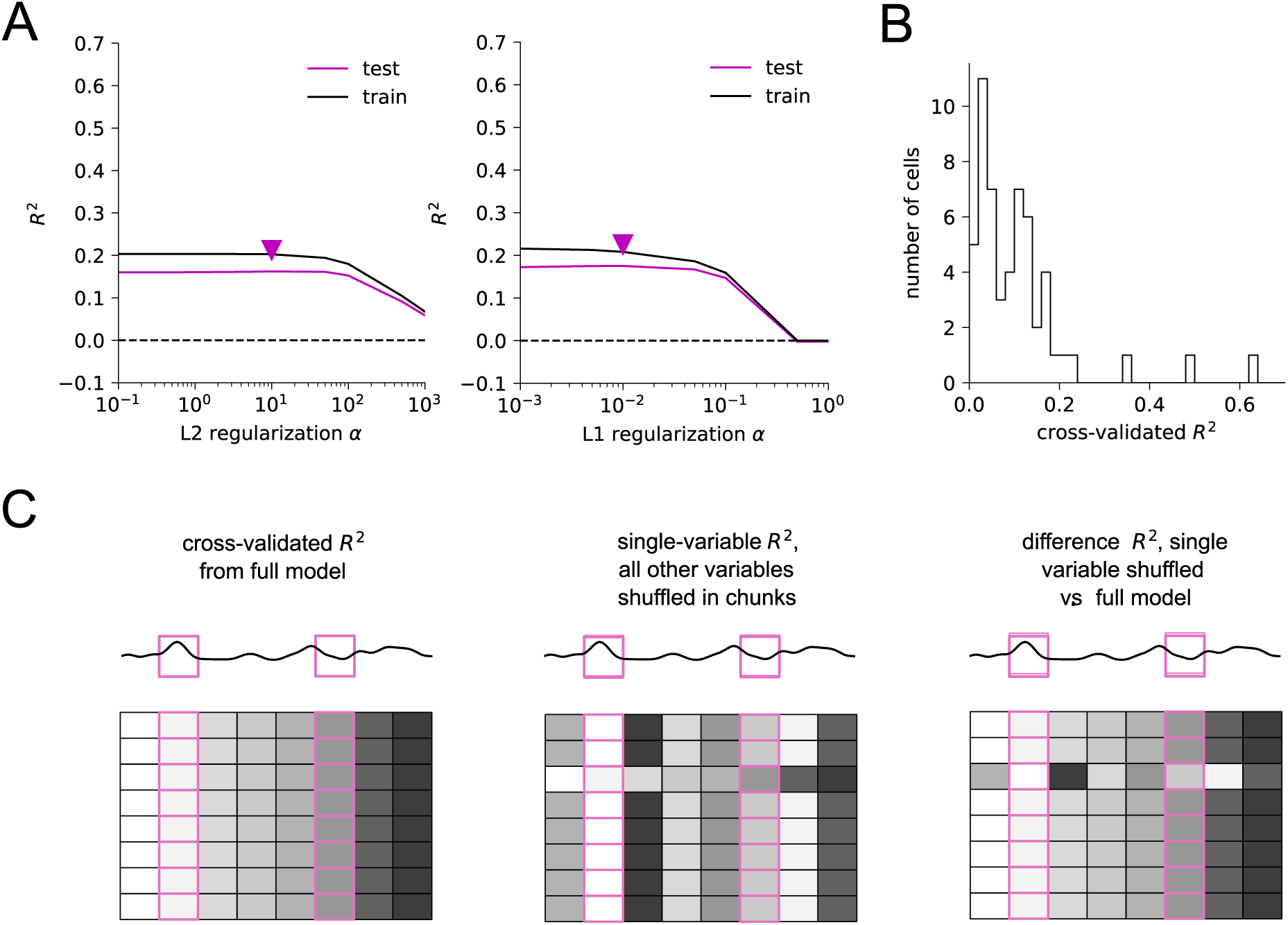
GLM analysis approach and *R*^2^ quantifications. (**A**) Illustration of how the regularization weight *α* is determined for both, the L2 and the L1-regularized regressions. The change of train and test *R*^2^ is shown as function of the regularization weight *α*, for an example regression. We used the largest value of *α* before a considerable drop in *R*^2^ as indicated by the arrow. (**B**) Histogram of the cross-validated *R*^2^s for the full model with 8 regressors (swing/stance onset for FL, FR, HL, and HR) after the second L2-regularized regression for all cells. (**C**) Cross-validated *R*^2^ from the full model (left), from single-variable model (middle), and from a leave-one-variable-out model (right). In all models, we cut time series into decorrelated chunks of 200 ms bins (Methods), trained the GLM on 80% of chunks, and assessed the *R*^2^ on the left-out 20% of chunks. For single-variable models, we shuffled the chunks of all other variables before training to keep only the relationship between firing rates and the variable of interest. To assess the unique contribution of the variable of interest, we calculated the difference in *R*^2^ after shuffling the variable of interest vs. the *R*^2^ in the full model. Significance was determined based on 95% confidence intervals from 100 repeated shuffles.

**Figure A.12:**
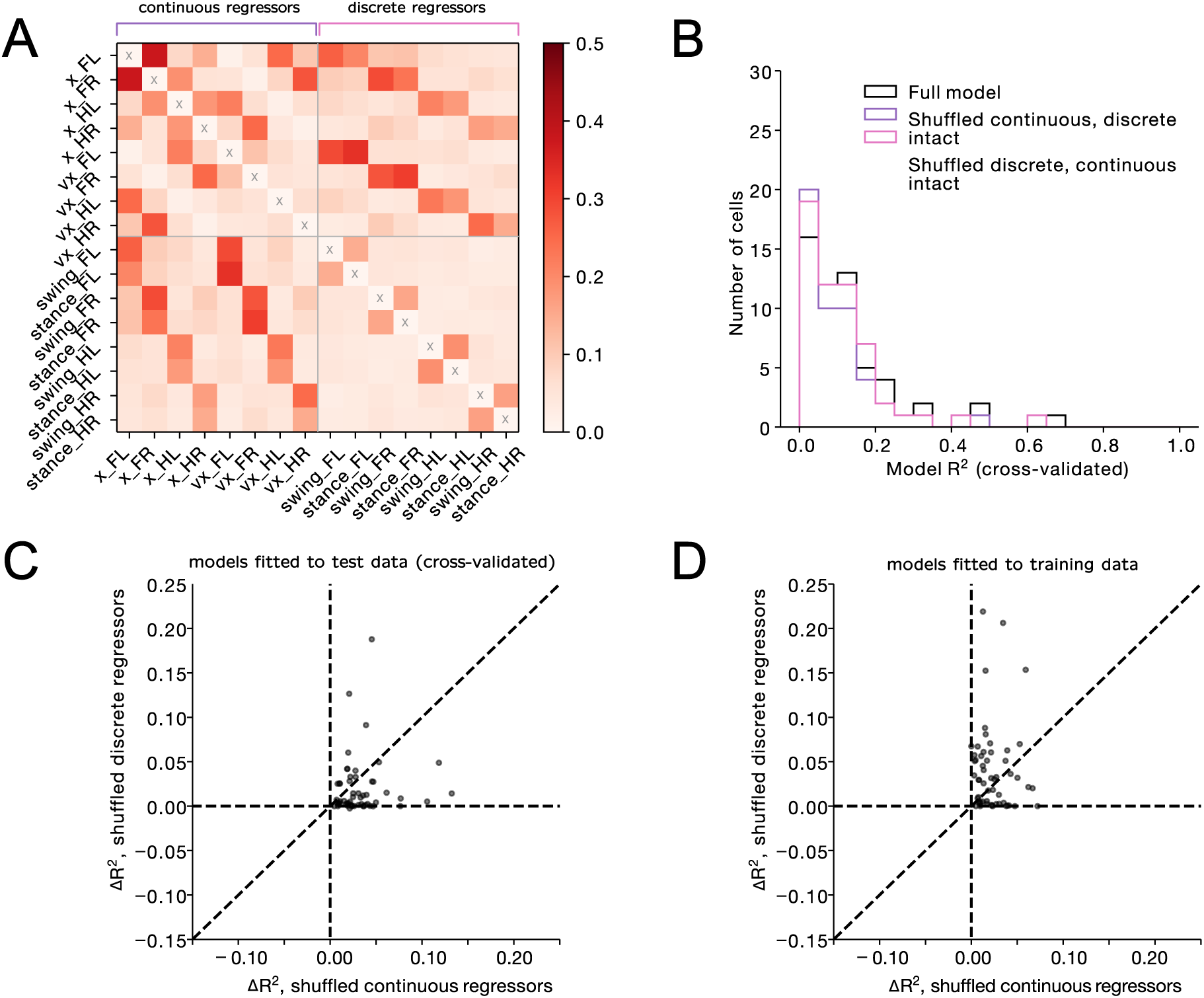
Δ*R*^2^ analysis : comparison between continuous and discrete regressor models. (**A**) Average (across animals and sessions) absolute Pearson’s *r* between different regressors reveals correlation structure in the regressors, showing that discrete swing and stance event kernels (swing_FL,FR,HL,HR and stance_FL,FR,HL,HR) are correlated with continuous variables (x position and speed along x). Autocorrelations not shown for better visibility of cross-correlations. (**B**) Comparison of the distribution of cross-validated *R*^2^ between regression models: Black, a model containing 8 event regressors (swing/stance onset for FL, FR, HL, and HR) as well as paw-specific continuous variables, i.e., the four paw x-positions and speeds. Purple, a model with shuffled continuous regressors, i.e. containing only event-related information. Pink, a model with shuffled discrete regressors, i.e. containing only kinematic variables. Each data point is the cross-validated *R*^2^ value of a different MLI for the discrete vs. continuous regression model, where the regression model was fitted on all trials from a single session. (**C**) Difference in *R*^2^ between the full model (black trace in the histogram in B) and the respective reduced model with paired cross-validated *R*^2^ values of a different MLI for the discrete vs. continuous regression model. (**D**) Same depiction as in C but for non-cross-validated *R*^2^ values, *i.e.*, *R*^2^ calculated on the training set.

**Figure A.13:**
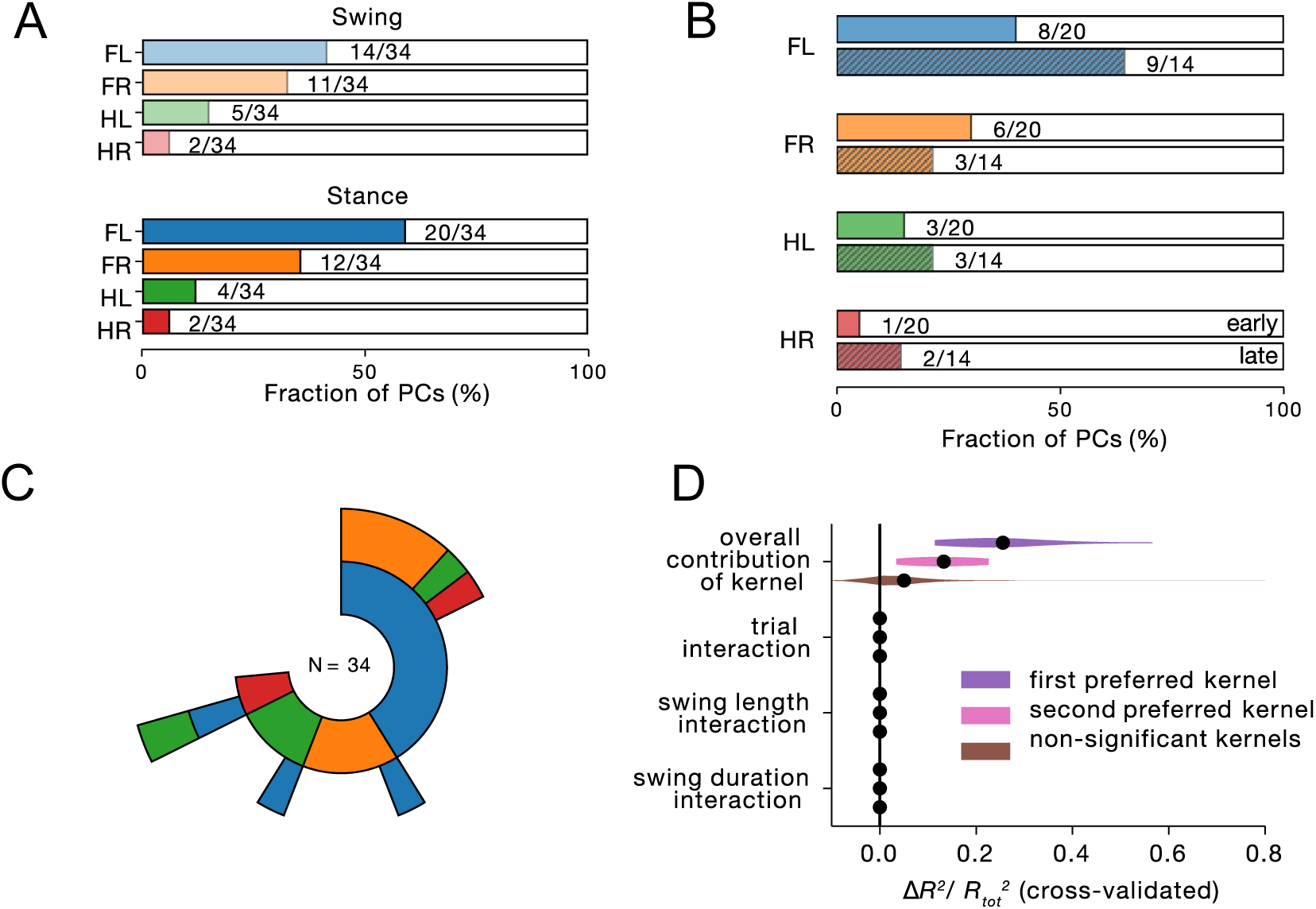
GLM analysis disentangles unique PC contributions of correlated behavioral events. Similar depection as Fig. 7 but for PCs. (**A**) Fraction of cells with significant single-variable *R*^2^ for different event types and paws. (**B**) Fraction of cells with significant difference in *R*^2^ (pooling swing and stance regressors) in early (upper, solid bars) and late (lower, striped bars) sessions (see Methods for details). (**C**) Inner circle: 25 out of 34 cells had at least one significant event regressor (swing or stance, difference in *R*^2^), color-coded by paw identity. Second circle, 9 out of 34 cells had a significant second event regressor. Third circle, 1 out of 34 cells had a significant third event regressor. (**D**) Unique contribution of different regressors (difference in *R*^2^) to overall explained variance 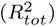. Purple, overall and interaction terms for first preferred event regressor (determined by absolute weight); pink, second preferred regressor; brown, non-significant event regressors.

## Notes

### Competing Interest Statement

The authors have declared no competing interest.

### Summary of Updates

We added an extensive misstep analysis to the manuscript, which proved to be extremely informative. This analysis allowed us to quantify corrective movements and activity at a fine-grained resolution and revealed important insights into how cerebellar activity relates to locomotor errors and adaptation. Importantly, this analysis could only be performed using the LocoReach task, which leverages the full potential of its implementation to enable high-resolution, paw-specific behavioral tracking during complex locomotion and link it to single-cell activity. We believe this aspect represents a key strength of the study and, together with careful behavioral quantifications, optogenetics, GLM analysis.

